# Scg2 drives reorganization of the corticospinal circuit with spinal premotor interneurons to recover motor function after stroke

**DOI:** 10.1101/2025.01.21.634186

**Authors:** Tokiharu Sato, Yuka Nakamura, Kana Hoshina, Ken-ichi Inoue, Masahiko Takada, Masato Yano, Hitoshi Matsuzawa, Masaki Ueno

**Affiliations:** Department of System Pathology for Neurological Disorders, Brain Research Institute, Niigata University, Niigata, Niigata, 951-8585, Japan; Center for the Evolutionary Origins of Human Behavior, Kyoto University, Inuyama, Aichi 484-8506, Japan; Department of Neurology, Osaka University Graduate School of Medicine, Suita, Osaka 565-0871, Japan; Division of Neurobiology and Anatomy, Graduate School of Medical and Dental Sciences, Niigata University, Niigata, Niigata, 951-8585, Japan; Center for Integrated Human Brain Science, Niigata University, Niigata, 951-8585, Japan; Center for Advanced Medicine and Clinical Research, Kashiwaba Neurosurgical Hospital, Sapporo, 062-8513, Japan

## Abstract

Brain injuries such as stroke damage neural circuitry and lead to functional deficits. Spared motor pathways are often reorganized and contribute to functional recovery; however, the connectivity and molecular mechanisms that drive the reorganization are largely unknown. Here, we demonstrate structural and functional connectivity reformed by spared corticospinal axons after stroke and determine a key secretory protein that drives the reorganization. We first found that corticospinal axons innervate specific areas of the denervated cervical cord after stroke. Anatomical and photometric analyses reveal that the axons reconnect to premotor V2a interneurons. Kinematic analyses of forelimb movements and chemogenetic silencing reveal their contribution to motor recovery. Translated mRNA expression analyses of V2a interneurons and astrocytes in the denervated cervical cord reveal diverse transcripts upregulated in the rewiring process. In particular, a secretory protein Scg2 is upregulated by injury-induced purinergic ATP signals and rehabilitative training-induced neural activity and possesses an ability to promote axon growth via cAMP and S6 signaling. Scg2 overexpression in the denervated cervical cord enhances axon rewiring, while the Scg2 knockdown attenuates it. The present data reveal the neural substrate and molecular mechanism essential to induce reorganization and recovery of the motor system, providing fundamental therapeutic targets for CNS injuries.

## Introduction

Stroke is one of the leading causes of neurological deficits resulting in motor and sensory dysfunction (Murphy and Corbett, 2009; Joy and Carmichael, 2021). While pharmacological and mechanical treatments are effective for reperfusion in the acute phase, therapeutic options for recovery are limited in the chronic phase except for rehabilitation (Campos et al., 2023; Tsivgoulis et al., 2023). The loss of neurons leads to disconnection of diverse intrinsic circuits that connect local and distal areas, resulting in functional impairments. Notably, however, patients often exhibit a limited but significant recovery over time. Previous studies have demonstrated that plastic changes in the spared neural networks correlate well with the process of recovery in both human and animal models (Cramer, 2008; Murphy and Corbett, 2009; Nudo, 2013; Carmichael et al., 2017; Ward, 2017; Isa et al., 2019). This raises the possibility that promoting the process of reorganization would be a promising therapeutic strategy. However, the precise targets of neural substrates and molecular mechanisms that drive the reorganization still remain largely unknown.

Stroke often damages the motor pathways such as the corticospinal tract (CST), the descending pathway for voluntary and skilled movement sending commands from the cerebral cortex to the spinal cord (Poter, Robert, 1995; Lemon, 2008; Boyd et al., 2017). In animal models, spared CST axons are often rewired to form compensatory circuits after injuries (Isa et al., 2019; Liu et al., 2021). For example, the axons regrow in the denervated area of the spinal cord and contribute to the recovery (Lee et al., 2004; Ueno et al., 2012; Lindau et al., 2014; Wahl et al., 2014; Morecraft et al., 2016; Okabe et al., 2016; Kaiser et al., 2019). Ipsi- or contralesional CST axons are rewired depending on the location and size of the stroke in mice and monkeys (Morecraft et al., 2016; Sato et al., 2021). These suggest that reorganization of the corticospinal circuitry would be essential for motor recovery. However, neural and molecular targets to control the reorganization remain to be determined. Specifically, what kind of connections are reformed and what molecular mechanisms control the axon rewiring are largely unknown. The CST comprises multiple projections from the motor and sensory cortical areas, with diverse connections within the spinal cord (Asante and Martin, 2013; Kamiyama et al., 2015; Wang et al., 2017; Ueno et al., 2018; Usuda et al., 2022). For example, we have shown that the axons from the motor cortex project to intermediate to ventral laminae of the spinal cord and primarily connect with motor-related spinal interneurons such as Chx10^+^ V2a interneurons in mice (Ueno et al., 2018). In contrast, the axons from the sensory cortex project dorsally and connect to interneurons involved in sensory functions. Each circuit regulates specific aspects of sensorimotor functions in behaviors (Liu et al., 2018; Ueno et al., 2018). After injuries, spared axons regrow and connect to segmental interneurons and propriospinal neurons that would compensate for the lost motor functions in rodents (Bareyre et al., 2004; Ueno et al., 2012). However, detailed connections of the compensatory circuits contributing to recovery are not comprehensively explored, in contrast to the detailed connection map in intact corticospinal circuits (Wang et al., 2017; Ueno et al., 2018; Carmona et al., 2024). Especially, the types of spinal interneuron targeted by the innervating CST axons as well as their functional contribution have not been elucidated yet.

Molecular mechanisms that control the axon rewiring are also not fully understood. Neural connections, especially in denervated areas, are reorganized in sequential steps after the injuries (Liu and Chambers, 1958; Raisman, 1969; Tsukahara, 1981). Injuries cause degeneration of neurons and axons extending to target areas, where they lead to disconnection from target neurons and a reaction of astrocytes and microglia. Spared axons then begin to sprout and grow to appropriate target neurons and reform synaptic connections, while some are eliminated for refinement. Certain molecular mechanisms in growing axons and surrounding cells within the target area will cooperatively facilitate the process of reorganization. In this context, previous studies had explored the molecules for the axon regrowth by focusing on factors putatively secreted in the denervated area. Expression analyses of candidate genes known for axon growth revealed that BDNF and CNTF could induce axon innervation of the CST (Ueno et al., 2012; Jin et al., 2015). Transcriptomic analyses using microarrays or RNA-seq in different injury contexts further demonstrated diverse gene expressions in the denervated areas (Bareyre et al., 2002; Maier et al., 2008; Li et al., 2010; Kaiser et al., 2019; Tsujioka and Yamashita, 2019; Chang et al., 2022). However, essential molecules that induce axon rewiring have been limitedly identified except for GDF10 in the cerebral cortex (Li et al., 2010). These approaches might fail to reveal key molecules due to a lack of sensitivity to detect expression changes that occur in a cell type-specific manner. Consequently, the cellular and molecular mechanisms that trigger the reorganization are not well understood.

In the present study, we investigated the neural connectivity reformed in the corticospinal circuits for recovery and further explored the molecular mechanisms that drive the rewiring in a mouse stroke model. We found that spared CST axons re-connect to a specific type of spinal interneuron, which were required for a specific aspect of recovery in forelimb movement. We further performed cell type-specific translated mRNA expression analysis and revealed that a secretory protein, secretogranin II (Scg2), was upregulated in astrocytes and target neurons by injury-induced purinergic signals and neural activity, respectively. Finally, we revealed that Scg2 is essential for the induction of axon reorganization. The present data demonstrate the neural and molecular bases of circuit reorganization induced after stroke, which could be important targets for developing therapeutic approaches for CNS injuries.

## Results

### Spared CST axons grow into the denervated side of the cervical cord after stroke

Spared CST axons are reorganized in the cervical spinal cord after cortical injuries (Ueno et al., 2012; Lindau et al., 2014; Kaiser et al., 2019; Sato et al., 2021). We especially showed that the axons from the contralesional cortex grew into the denervated side across the midline, when a large size of stroke was induced in the cortical area where the CST axons originated (rostral forelimb area [RFA], caudal forelimb area [CFA], and primary somatosensory cortex [S1]; **Fig. 1a**; Ueno et al., 2018; Sato et al., 2021). We first examined the spatiotemporal changes in the axon innervation in a large-sized stroke induced by a photothrombotic method, which degenerated most of the CST fibers derived from the lesion side of the cortex (**Fig. 1a**, **b**, **Supplementary** Fig. 1). Spared CST axons were labeled from the contralesional motor cortex with an anterograde tracer, biotinylated dextran amine (BDA), after stroke (**Fig. 1a**). CST axons began to increase in the denervated side of the cervical cord from day 7 and the density peaked at day 28–56 (Fig. 1c, d). There was a slight decrease on day 56 at the C6–7 level, consistent with previous reports showing axon pruning during the chronic phase (**Fig. 1d**) (Bareyre et al., 2004; Ueno et al., 2012; Kaiser et al., 2019). The number of midline-crossing axons also increased at day 7, while the number did not significantly change over time (**Fig. 1e**). These indicate that spared CST axons cross the midline early after stroke, followed by an increase of growth and branching in the denervated side with the CST fibers that already innervate in the intact spinal cord.

**Figure 1.**
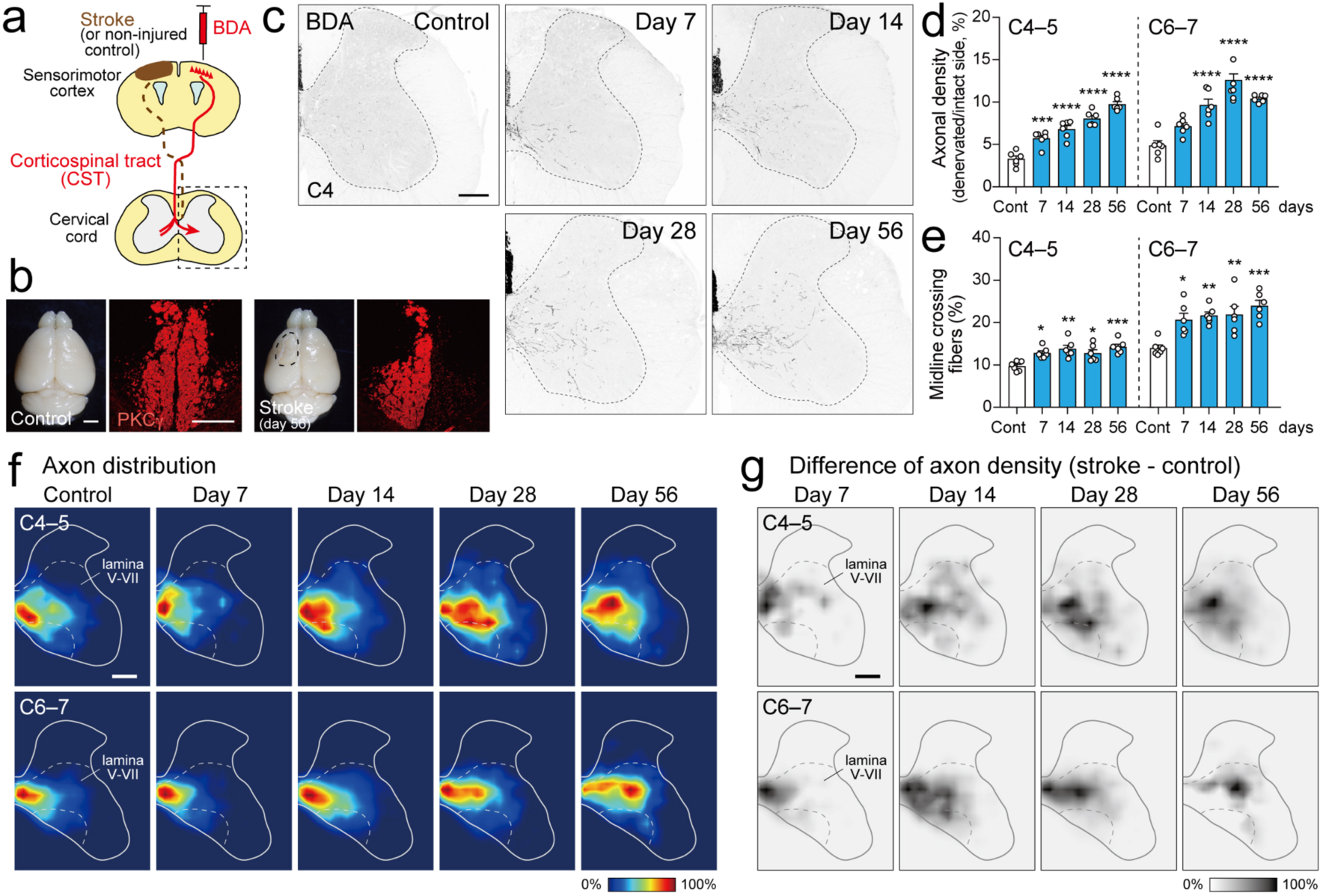
CST axons are rewired in the denervated side of the cervical cord after stroke. **a**, Schematic representations of anterograde labeling of CST from the contralesional motor cortex in a stroke model. **b**, Representative images of the dorsal view of the brain and PKCψ-stained CST in the dorsoventral column of the cervical cord in control and stroke mice at day 56. The dotted area indicates the lesion. Note that most CST axons from the lesion side degenerate. **c**, Representative BDA-labeled CST axons in the denervated side of the cervical cord in control and at day 7–56 after stroke (C4 level). The dotted lines mark the boundary between the gray and white matter. **d**,**e**, Quantification of the axonal density in the denervated side (**d**) and the midline crossing CST fibers (**e**) at C4–5 and C6–7 levels post-stroke. N = 6 mice, one-way ANOVA followed by Tukey’s test, \**p* < 0.05, \*\**p* < 0.01, \*\*\**p* < 0.001, and \*\*\*\**p* < 0.0001. **f**,**g**, Heatmaps showing the CST axon distribution in the denervated side at C4–5 and C6–7 post-stroke (**f**) and the difference in axon density (subtraction of controls from the stroke groups; **g**). Scale bars, 2 mm (left panel of **b**); 100 µm (right panel of **b**); 200 µm (**c**, **f**, **g**).

We next analyzed the pattern of axon innervation in heatmaps (**Fig. 1f**). The axons mainly projected to the medioventral area including laminae V–VIII, which was similar with the distribution in non-injured control mice. However, they showed a tendency to project more dorsolateral and ventrolateral areas within the laminae over time (**Fig. 1f, g**). The data suggest that the spared CST axons innervate specific areas of the spinal gray matter after stroke.

### Rewired CST axons connect to V2a interneurons after stroke

The innervation in specific spinal areas implied formation of certain specific connections. We next asked to what target neurons the CST axons connect to reorganize the circuit.

We previously showed that rewired CST axons connected to segmental and propriospinal neurons after cortical injury (Ueno et al., 2012). However, the precise neuronal subtype especially regarding the diverse spinal interneuron subtypes classified along the dorsoventral axis during the development remained unknown (**Fig. 2a**; Goulding, 2009; Alaynick et al., 2011). We first compared the spatial distribution of reorganized CST axon projections and spinal interneuron subtypes, which were labeled with Cre and GFP reporter mice (Ueno et al., 2018) (**Fig. 2b**, **Supplementary** Fig. 2a, b). Correlation coefficient analyses revealed a high spatial correlation of innervating CST axons with the locations of Chx10^+^ (V2a), Dbx1^+^ (V0/dl6), and Olig3^+^ (V1–3) interneurons in addition to Nkx2.2^+^ (V3) and Isl1^+^ (dl3) interneurons in C4–5 and C6–7, respectively (**Fig. 2c**, **Supplementary** Fig. 2c, d). In contrast, the locations of dorsal populations such as Vglut3^+^ (D), Ptf1a^+^ (dl4-dlLa), and Lmx1b^+^ (dlL-dl5) interneurons showed a lower correlation (**Fig. 2a–c**, **Supplementary** Fig. 2), which are originally targeted by CST axons from the sensory cortex (Ueno et al., 2018).

**Figure 2.**
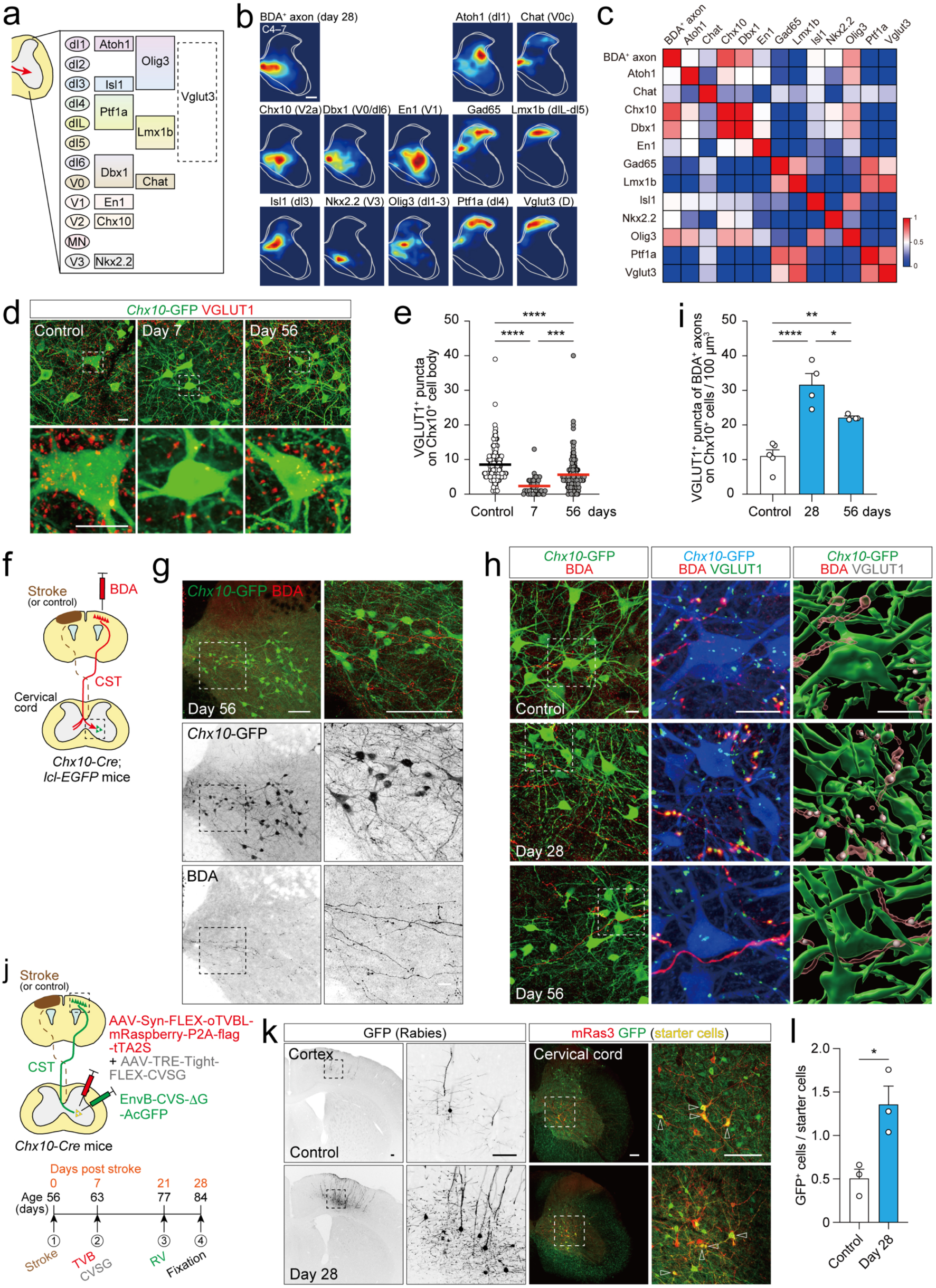
CST axons anatomically connect with V2a interneurons after stroke. **a**, Schematic representations of spinal interneuron subtypes with marker genes. **b**, Heatmaps showing the distribution of BDA^+^ CST axon projections at day 28 post stroke and GFP-labeled spinal interneurons with Cre-reporter mice at C4–7. **c**, Correlation coefficient matrix of the location of CST axons and spinal interneurons. **d**, Representative confocal images of VGLUT1^+^ excitatory synaptic contacts (red) onto GFP^+^ V2a interneurons (green) in control and stroke *Chx10-Cre*;*lcl- EGFP* mice at day 7 and 56. Lower panels show high magnifications of the dotted areas. **e**, Quantification of the number of VGLUT1^+^ puncta on *Chx10*-GFP^+^ cell body. N = 153 (control), 41 (day 7) and 114 (day 56) cells, one-way ANOVA followed by Tukey’s test, \*\*\**p* < 0.001 and \*\*\*\**p* < 0.0001. **f**, Schematic representations of BDA injection into the contralesional motor cortex of *Chx10*-*Cre*;*lcl*-*EGFP* mice. **g**, Representative images of BDA^+^ CST axon innervation (red) to the location of *Chx10*-GFP^+^ interneurons (green) post-stroke. Right panels show higher magnification views of the dotted areas in the left panels. **h**, Representative images of CST axon connections onto V2a interneurons post-stroke. Left panels, confocal images of BDA^+^ CST axons (red) and *Chx10*-EGFP*^+^* interneurons (green). Middle, higher magnification z-stack images of the dotted areas in the left panels. BDA (red), *Chx10*-EGFP (blue), and VGLUT1 (green). Right, reconstructed IMARIS images of the middle panels. BDA (red), *Chx10*-EGFP (green), and VGLUT1 (gray). **i**, The number of synaptic contacts of CST axons onto V2a interneurons. N = 5 (control) and 4 (day 28 and 56 post stroke) mice, one-way ANOVA followed by Tukey’s test, \**p* < 0.05, \*\**p* < 0.01, \*\*\*\**p* < 0.0001. **j**, Experimental diagram of monosynaptic retrograde tracing from V2a interneurons with rabies virus. Bottom, experimental timeline. **k**, Representative images of the AcGFP^+^ traced neurons in the motor cortex ipsilateral to the injection side (black, left panels) and the mRaspberry3^+^/AcGFP^+^ starter cells in the cervical cord (red and green, arrowheads, right panels) in control and stroke mice at day 28. Right panels: higher magnification views of the dotted areas are presented in the right side. **l**, The number of AcGFP^+^ corticospinal neurons mono-synaptically and retrogradely traced from V2a interneurons. N = 3 mice, Student’s *t*-test, \**p* < 0.05. Scale bars, 200 µm (**b**); 100 µm (left of **g**, **k**); 50 µm (right of **g**); 20 µm (**d**, **h**).

Among the candidate target neurons, we focused on Chx10^+^ V2a neurons, which have been reported as the cardinal premotor interneurons involved in motor output (Crone et al., 2008; Azim et al., 2014b; Ueno et al., 2018). We first examined whether synaptic connections were changed in V2a neurons after stroke. The number of VGLUT1^+^ excitatory presynaptic contacts, which CST axon terminals possess (Maier et al., 2008; Ueno and Yamashita, 2011), on GFP-labeled V2a neurons was significantly decreased at day 7 post-stroke in *Chx10*-*Cre*;*lox*-*CAT*-*lox*-*EGFP* (*lcl*-*EGFP*) mice, but then the number was found to recover at day 56 (**Fig. 2d**, **e**). We then examined synaptic contacts of reorganized CST axons onto V2a neurons. As predicted in the correlation coefficient analyses, CST axons innervated the areas where GFP^+^ V2a neurons were located (**Fig. 2f**, **g**). We further found that the axon terminals labeled by BDA and VGLUT1 had contacts onto GFP^+^ V2a neurons and the number significantly increased at days 28–56 after stroke (**Fig. 2f–i**). The number peaked at day 28 and decreased at day 56, correlating with the pruning process of the axons. We further evaluated their connections by performing monosynaptic retrograde tracing from V2a neurons using EnvB-coated G-deleted rabies virus of CVS strain (**Fig. 2j**). We confirmed that this strain had an efficient ability to move retrogradely through the long descending pathway to label layer V neurons, which could not be achieved by using the widely used SADB19 strain (Ueno et al., 2018). The number of retrogradely labeled neurons in the motor cortex monosynaptically connected with V2a neurons was significantly increased at day 28 after stroke (**Fig. 2k**, **l**). Collectively, the data indicate that the reorganized CST axons target V2a neurons to reconnect the circuit.

To determine whether the CST axons formed functional connections with V2a neurons, we further examined physiological responses of V2a neurons to the stimulation of CST axons by using a fiber photometry and optical stimulation. Intracellular Ca^2+^ responses were recorded with jGCaMP7s expressed in V2a neurons on the denervated side by injecting AAV-Syn-DIO-jGCaMP7s into the cervical cords of *Chx10*-*Cre* mice (**Fig. 3a**, **b**). ChrimsonR, a red-shifted variant of ChR2, was further introduced in the CST axons by injecting adeno-associated virus (AAV) into the contralesional motor cortex (**Fig. 3a**, **b**, **Supplementary** Fig. 3a). Calcium responses were then recorded upon light-stimulated activation of CST axons. In non-injured mice, single-pulse stimulation caused a slight increase in jGCaMP7s signals, indicating synaptic connections between midline-crossing CST axons and V2a neurons even in controls (**Fig. 3c–e**). We then found that the calcium signals dramatically increased at day 28 and 56 post-stroke (**Fig. 3c–e**). The signals slightly decreased on day 56 compared to day 28, mirroring the pruning process of the anatomical connections (**Fig. 2h**, **i**). Taken together, the results indicate that spared CST axons form a refined functional circuit with V2a neurons after stroke.

**Figure 3.**
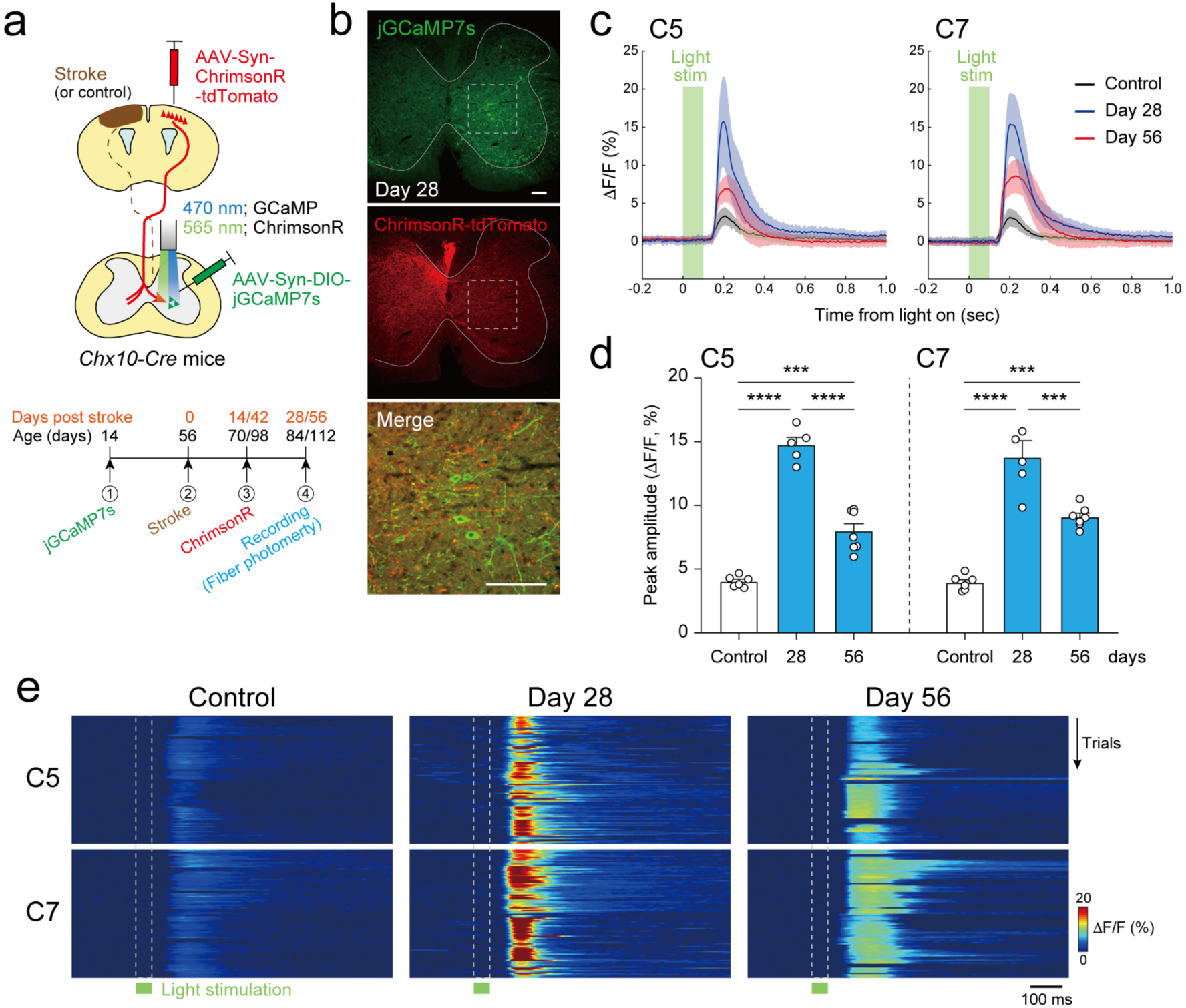
Functional connectivity of CST axons and V2a interneurons after stroke. **a**, Experimental diagram of fiber photometry and optogenetic stimulation. ChrimsonR-expressing CST axons were optically stimulated and calcium responses of jGCaMP7s-expressing *Chx10^+^* interneurons were recorded at C5 and C7. Bottom, experimental timeline. **b**, Representative images of jGCaMP7s (green) and ChrimsonR-tdTomato expression (red) in the cervical cord at day 28 post-stroke. Bottom, a magnified view of the dotted area. Scale bars, 100 µm. **c**, Representative optically-evoked calcium responses of V2a interneurons at C5 (left) and C7 (right) levels in control (black) and stroke mice at day 28 (blue) and 56 (red). The solid lines and shaded areas represent the average and standard deviation of fluorescence changes, respectively. **d**, Peak amplitude of calcium responses. N = 6, 5 and 7 mice for control, day 28, and day 56, respectively; one-way ANOVA followed by Tukey’s test, \*\*\**p* < 0.001 and \*\*\*\**p* < 0.0001. **e**, Representative kymographs of calcium responses evoked by optical stimulation at C5 and C7 in controls and at day 28 and 56 post-stroke (50 trials/one mouse). The green bars and the dotted lines indicate the timing of photostimulation.

### V2a interneurons are required for motor recovery after stroke

Previous studies revealed that contralesional CST axons contribute to spontaneous motor recovery in a large size of cortical injury (Ueno et al., 2012; Wahl et al., 2014). We further asked whether the target V2a neurons that were connected with rewired CST axons were involved in the recovery process. We first examined motor deficits and subsequent recovery process of skilled forelimb movement by using a single pellet reaching task, and further interrupted the activity of V2a neurons by using a chemogenetic ion channel receptor PSAM^4^-GlyR (Magnus et al., 2019). We established an experimental system for motion analyses that automatically traced the trajectories of the reaching arms in two diagonally recorded movies by DeepLabCut (Mathis et al., 2018) and enabled to export the coordinate data into the motion analyses software, KinemaTracer (Ueno and Yamashita, 2011; Ueno et al., 2018; Nakamura et al., 2023) (**Fig. 4a**).

**Figure 4.**
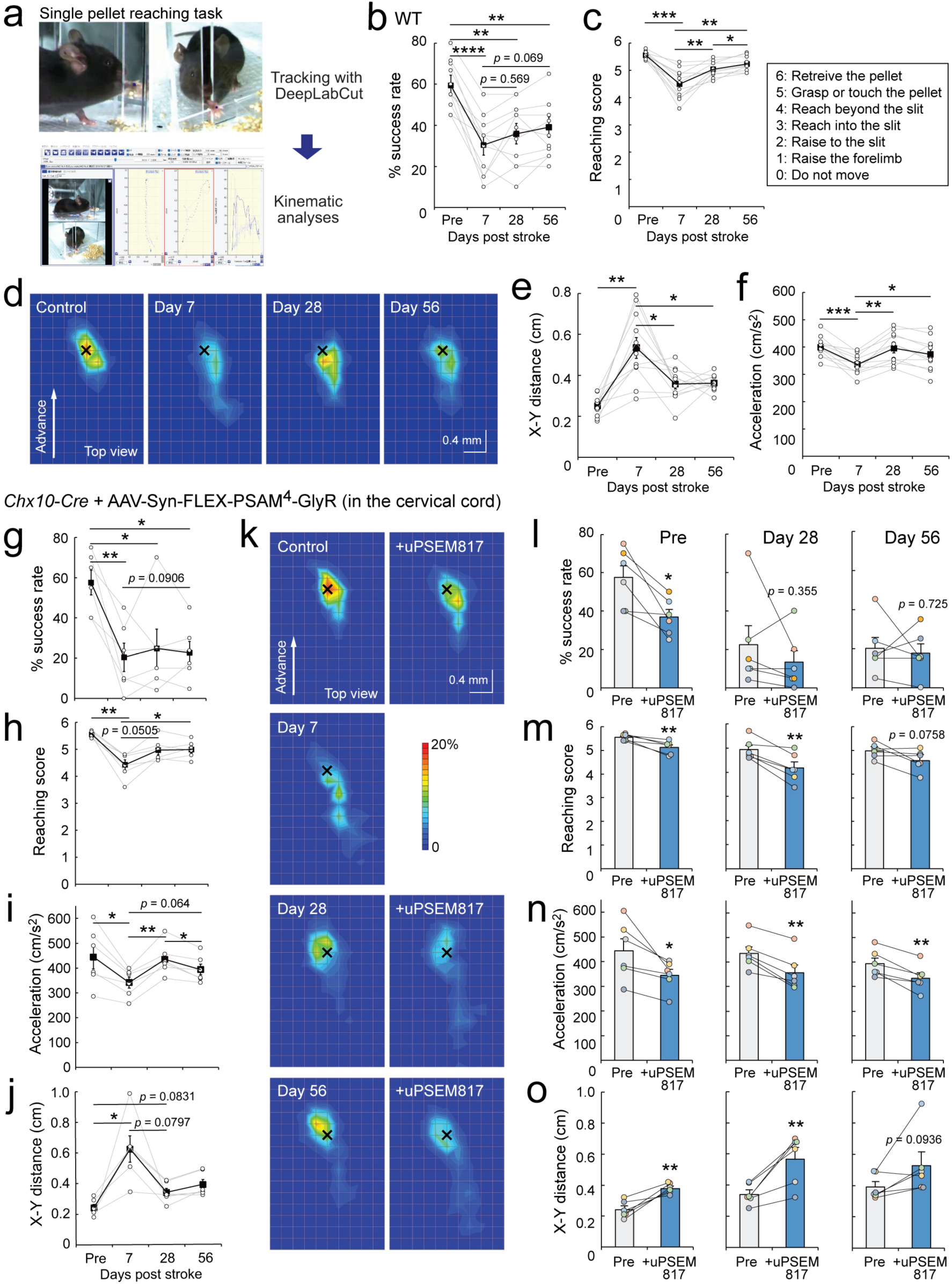
V2a interneurons contribute to the recovery of skilled motor functions after stroke. **a**, Experimental system to analyze the forelimb movements in the single pellet reaching task by automatically tracing paw movements with DeepLabCut followed by kinematic analyses in KinemaTracer. **b**,**c**, Success rate (**b**) and reaching score (**c**) pre- and post-stroke in wild-type (WT) mice. Reaching score was defined as the criteria in **c**. N = 11 mice, one-way ANOVA followed by Tukey’s test, \**p* < 0.05, \*\**p* < 0.01, \*\*\**p* < 0.001, \*\*\*\**p* < 0.0001. **d**, Heatmaps showing the probabilities of far distal reaching positions of the paw relative to the pellet position (×) in the top view in control and post-stroke mice at days 7–56. **e**,**f**, Distance between the distal positions of the paw and the pellet in x-y axis (**e**) and the acceleration of the forelimb in the advancement step (**f**) pre- and post-stroke. N = 11 mice, one-way ANOVA followed by Tukey’s test, \**p* < 0.05, \*\**p* < 0.01, \*\*\**p* < 0.001. **g**–**j**, Recovery of motor functions after stroke in *Chx10-Cre* mice infused with AAV-Syn-FLEX-PSAM^4^-GlyR in the denervated side of the cervical cord. Success rate (**g**), reaching score (**h**), acceleration of the forelimb in the advancement step (**i**) and the distance between the distal positions of the paw and the pellet in x-y axis (**j**). N = 6 mice, one-way ANOVA followed by Tukey’s test, \**p* < 0.05, \*\**p* < 0.01. **k**, Heatmaps showing the probabilities of the far distal reaching positions of the paw relative to the pellet position (×) in PSAM^4^-GlyR-expressing *Chx10-Cre* mice before and after uPSEM817 treatment pre- and post-stroke. **l**–**o**, Impairment of recovered functions after silencing PSAM^4^-GlyR expressing V2a neurons with uPSEM817 treatment. Success rate (**l**), reaching score (**m**), acceleration of the forelimb (**n**), and distance between the positions of the paw and the pellet (**o**) pre- and post-uPSEM817 treatment in controls and day 28 and 56 post-stroke. N = 6 mice, paired *t*-test, \**p* < 0.05, \*\**p* < 0.01.

Success rates in retrieving pellets were severely affected at day 7 and then showed a tendency of partial recovery but not statistically significant (**Fig. 4b**). Since the success rate values were variable and would not be suitable to evaluate qualitative improvement as previously discussed (Lemke et al., 2019), we further evaluated a reaching score that subdivided the quality of reaching in 7 rating points (Ueno et al., 2018). The scores worsened at day 7 but then recovered spontaneously at day 28 and 56 (**Fig. 4c**, **Supplementary Video 1**). Kinematic analyses further revealed that the distal positions of the paw during the reaching were disrupted at day 7 but then recovered at day 28 and 56 (**Fig. 4d**, **e**, **Supplementary** Fig. 4b). The acceleration of the reaching (cm/s^2^) was also impaired at day 7 and then recovered at day 28 and 56 after the stroke (**Fig. 4f**).

We next induced PSAM^4^-GlyR expression in V2a neurons by infusing AAV-Syn-FLEX-PSAM^4^-GlyR into the denervated side of the cervical cord of *Chx10*-*Cre* mice (**Supplementary** Fig. 4a). The reaching score, final reaching positions on the XY axis, and acceleration of the reaching spontaneously recovered after the stroke (**Fig. 4g–j**). However, the scores worsened when the Chx10^+^ V2a neurons were silenced by treatment with PSAM^4^-GlyR ligand uPSEM817 at day 28 and 56 post-stroke (**Fig. 4k–o**, **Supplementary** Fig. 4c, **Supplementary Video 2**). These results indicate that V2a interneurons are required for spontaneous motor recovery after stroke.

### Cell-type specific translated mRNA expressions in the denervated cervical cord after stroke

The molecular mechanisms underlying the induction of CST reorganization remain unknown. We sought to find key molecules for reorganization that might be secreted by the cells in the denervated target area to induce growth and rewiring of spared axons. Transcriptome analysis of bulk spinal tissues could not identify crucial molecules involved in the rewiring (Bareyre et al., 2002; Maier et al., 2008; Li et al., 2010; Kaiser et al., 2019; Tsujioka and Yamashita, 2019), which might miss to detect subtle yet important changes of gene expressions in specific cell types. Thus, we leveraged a RiboTag method that enables to detect cell-type specific expressions of translated mRNAs (‘translatome’) (Sanz et al., 2009), which correlate more precisely to the actually expressed proteins than the data in transcriptomic analyses (Schwanhäusser et al., 2011). We targeted translated mRNAs of V2a neurons and neighboring astrocytes and microglia in the denervated side, which would react to axon degeneration and might engage in circuit or tissue repair supporting axonal elongation and synaptic formation (Yu et al., 2021; Hemati-Gourabi et al., 2022). *Chx10-Cre*, *Aldh1l1-CreER* (for astrocytes) and *Cx3cr1-CreER* mice (for microglia) were crossed with *RiboTag* mice that express HA-tagged ribosomal protein *Rpl22* in a Cre dependent manner, which allow to extract HA-tagged ribosome-bound mRNAs in Cre-expressing cells through immunoprecipitation (**Fig. 5a, b**). As the reorganization started at day 7 (**Fig. 1c–e**), we collected cell type-specific mRNAs in the denervated side on day 3, 7, 14, and 28. The specificity of HA-immunoprecipitated mRNAs was confirmed by the expression of cell-type specific marker genes in quantitative PCR and RNA-seq (**Fig. 5c**, **Supplementary** Fig. 5a) (Zhang et al., 2014). Principal component analysis (PCA) further revealed similarities in gene expression in each cell type and time point group (**Supplementary** Fig. 5b). We next examined differentially expressed genes (DEGs) in the stroke groups compared to controls in each cell type. We found that hundreds of transcripts were temporally up- or downregulated in V2a neurons, astrocytes, and microglia in the denervated side (**Fig. 5d**, **Supplementary** Fig. 5c).

**Figure 5.**
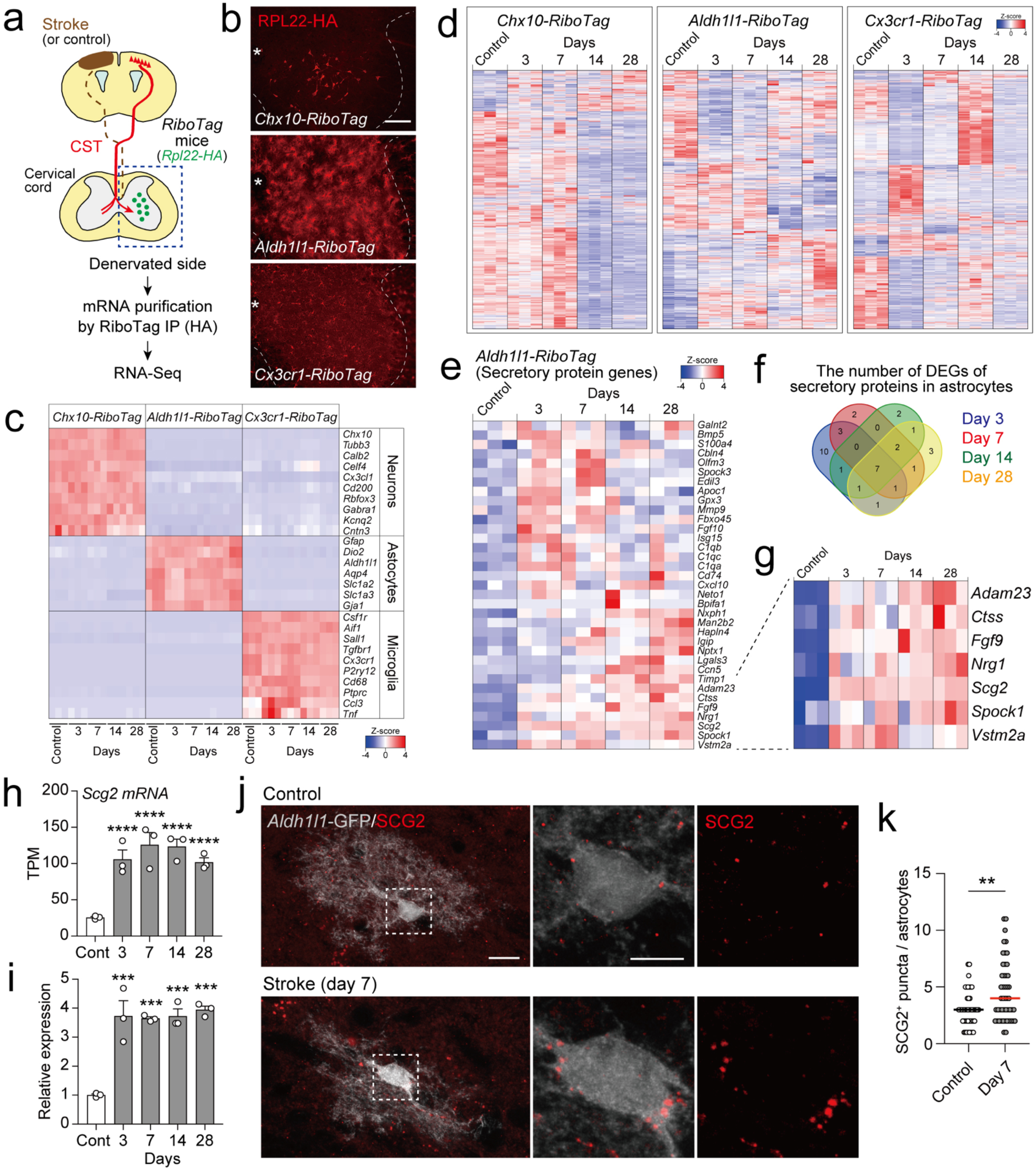
Cell-type specific translated mRNA expression analyses reveal *Scg2* upregulation in astrocytes after stroke. **a**, Experimental diagram of cell-type specific mRNA preparation and RNA-seq with the RiboTag method. **b**, HA-tagged RPL22 expression in the cervical cord of *Chx10-Cre*;*RiboTag*, *Aldh1l1-CreER*;*RiboTag* and *Cx3cr1-CreER*;*RiboTag* mice at the day 7 post-stroke. Asterisks represent the central canal. **c**, Heatmap showing expression of marker genes of neurons, astrocytes and microglia in RNA-seq data of each *RiboTag* mouse line. N = 3 mice for each group. **d**, Heatmaps showing the expression of DEGs in controls and at days 3–28 post- stroke in each *RiboTag* mouse line. **e**,**f**, Heatmap showing upregulated DEGs encoding secretory proteins in astrocytes (**e**) and their numbers in the Venn diagram (**f**). **g**, Heatmap showing seven constantly upregulated DEGs in astrocytes after stroke. **h**,**i**, Increased expression of *Scg2* mRNA after stroke in RNA-seq (**h**) and quantitative PCR (**i**). N = 3 for each condition. Multiple testing by Benjamini and Hochberg procedure (**h**), \*\*\*\**p* < 0.0001. One-way ANOVA followed by Tukey’s test (**i**), \*\*\**p* < 0.001. **j**, Representative confocal images of SCG2 expression (red) in *Aldh1l1*-GFP^+^ astrocytes (gray) at the denervated side of the cervical cord of control and stroke group at day 7 in *Aldh1l1-CreER*;*lcl-EGFP* mice. Middle and right panels show higher magnifications of the dotted areas. **k**, The number of SCG2^+^ puncta in *Aldh1l1*-GFP^+^ astrocytes. N = 46 (control) and 48 (day 7) cells from 2 and 3 mice, respectively; Student’s *t*-test, \*\**p* < 0.01. Scale bars, 100 µm (**b**); 10 µm (left in **j**); 5 µm (middle in **j**).

### *Scg2* is upregulated in astrocytes by ATP signals after stroke and promotes axon growth

Astrocytes engage in structural and functional support of synapses and formation of the extracellular matrix by secreting proteins that would support tissue and circuit repair (Dallérac et al., 2018; Hemati-Gourabi et al., 2022). We thus hypothesized that molecules secreted from astrocytes may induce the reorganization. We examined transcripts encoding secretory proteins among the upregulated DEGs in astrocytes and found total 35 genes at the tested time points after stroke (**Fig. 5e**, **f**). Based on the continuous increase in CST innervation (**Fig. 1c**, **d**), genes persistently upregulated might regulate the reorganization. Accordingly, we identified seven genes, *Adam23*, *Ctss*, *Fgf9*, *Nrg1*, *Scg2*, *Spock1*, and *Vstm2a* that were continuously upregulated from day 3 to 28 (**Fig. 5g**, **Supplementary** Fig. 5d). Among them, we focused on secretogranin II (*Scg2*), which showed the highest level of expression. SCG2 is known as a member of the chromogranin/secretogranin family regulating the secretory pathway that facilitate sorting and packaging of neuropeptides and hormones into dense-core secretory granules (Taupenot et al., 2003; Bartolomucci et al., 2011). We also noted that secreted SCG2 had been reported to act on neurons and induce neurite growth as well as engage in neuroprotection (Shyu et al., 2008; Kim et al., 2015). However, its physiological and pathological functions in injuries were largely unknown.

We first validated the upregulation of *Scg2* by quantitative PCR, showing an increase in *Scg2* mRNA expression in astrocytes, consistent with the RNA-seq data (**Fig. 5h**, **i**). We further performed immunohistochemical analyses of SCG2 in the cervical cord using *Aldh1l1-CreER*;*lcl-EGFP* mice to delineate the dendritic structures of astrocytes. SCG2 was observed as vesicle-like punctate signals, localized at the edges of the cell bodies and their processes (**Fig. 5j**). These signals likely correspond to secretory vesicles, as SCG2 is localized in the dense-core vesicles (DCVs) in cultured neuronal cells or astrocytes (Courel et al., 2010; Prada et al., 2011). We then found a significant increase in the number of SCG2^+^ puncta after stroke (**Fig. 5j**, **k**). These results indicate that *Scg2* expression is increased in the cervical astrocytes after stroke.

We next investigated whether SCG2 had an ability to promote axon growth in cultured neurons. Cerebellar granule neurons (CGNs) were cultured in dishes coated with axonal growth-inhibiting molecules, Nogo or chondroitin sulfate proteoglycans (CSPG), to mimic the inhibitory environment of the CNS (Atwal et al., 2008; Usher et al., 2010). Although they significantly inhibited neurite outgrowth, we found that SCG2 treatment enhanced the growth to control levels (**Fig. 6a**, **b**). We further sought the signaling molecules involved in the SCG2-induced neurite growth. Since *Scg2* knockout had inhibited ERK and PI3K/AKT pathways in zebrafish (Tao et al., 2018), we examined ERK and S6 activation, the downstream of PI3K/AKT. We found that SCG2 significantly increased the phosphorylation of ERK and S6 (**Fig. 6c**, **d**). Since the receptor for SCG2 is suggested to be a G protein-coupled receptor (GPCR) while not identified (Mitchell et al., 2020), we further examined the cAMP signaling pathway, a downstream of Gs-coupled receptor and a mediator of axon growth (Cai et al., 2001). By using cAMP sensor cAMPinG1-NE (Yokoyama et al., 2024), we found that SCG2 treatment elevated cAMP levels in cultured neurons (**Fig. 6e**). Concomitant treatment with SCG2 and SQ22536, an adenylate cyclase inhibitor, perturbed ERK and S6 activation (**Fig. 6f**, **g**). In addition, SQ22536 inhibited the effect of neurite outgrowth by SCG2 (**Fig. 6h**, **i**). These results indicate that SCG2 enhances neurite outgrowth through the cAMP pathway.

**Figure 6.**
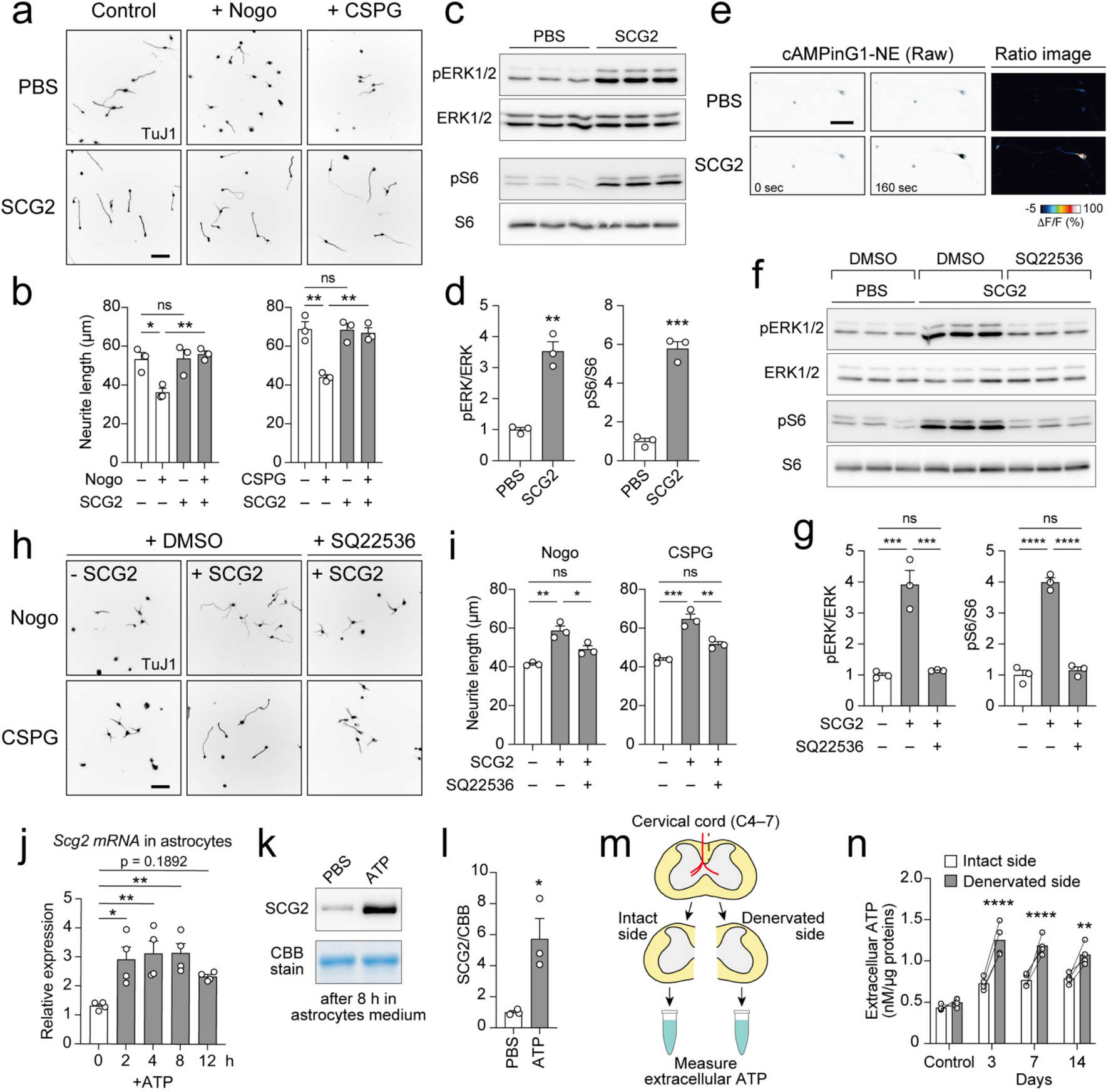
SCG2 promotes neurite outgrowth via cAMP signaling and its expression is induced by ATP stimulation. **a**, Representative images of neurite outgrowth of CGNs treated with or without SCG2 on control, Nogo and CSPG substrates. TuJ1 staining. **b**, Quantification of neurite length. N = 3, one-way ANOVA followed by Tukey’s test. **c**, Western blots showing phosphorylation of ERK and S6 proteins in CGNs 30 min after SCG2 treatment. Each lanes contains samples from three independent experiments. **d**, Quantitative data of pERK and pS6 immunoblots in **c**. N = 3, Student’s *t*-test. **e**, Raw fluorescence and ratio images of cAMPinG1- NE-expressing CGNs before (0 sec) and after PBS and SCG2 administration (160 sec). **f**, Western blots showing attenuated ERK and S6 phosphorylation by SQ22536 treatment in CGNs 30 min after SCG2 treatment. **g**, Quantitative data of pERK and pS6 in **f**. N = 3, one-way ANOVA followed by Tukey’s test. **h**, Representative images of neurite outgrowth of CGNs treated with or without SCG2 and SQ22536 on Nogo and CSPG substrates. TuJ1 staining. **i**, Quantification of neurite lengths of **h**. N = 3, one-way ANOVA followed by Tukey’s test. **j**–**l**, Expression of *Scg2* mRNA in cultured astrocytes (**j**) and secretion of SCG2 protein in conditioned media of astrocytes after ATP stimulation (**k**,**l**). N = 4, one-way ANOVA followed by Tukey’s test (**j**); N = 3, Student’s *t*-test (**l**). **m**, Schematic representations of measuring extracellular ATP concentration in the cervical cord after stroke. **n**, Quantitative data of extracellular ATP concentration in the control and stroke group at days 3–14. N = 4, two-way ANOVA followed by Sidak’s post hoc test. Scale bars, 50 µm. ns, not significant; \**p* < 0.05, \*\**p* < 0.01, \*\*\**p* < 0.001, \*\*\*\**p* < 0.0001.

We further examined the mechanism underlying the *Scg2* upregulation after stroke. Previous studies reported that the transcription factor REST negatively regulates *Scg2* expression (Prada et al., 2011; Kim et al., 2015). However, *Rest* expression was low and did not dramatically change in the RNA-seq data of astrocytes (data not shown). *Scg2* expression is also induced by Ca^2+^ influx and the cAMP/PKA/CREB signaling pathway (Iwase et al., 2014; Hasel et al., 2017). Since extracellular purines such as ATP are elevated in injured tissues (Wang et al., 2004; Davalos et al., 2005; Melani et al., 2005) and act through calcium channels and GPCRs, P2X and P2Y receptors, we tested whether ATP stimulation induces *Scg2* expression. ATP administration was found to significantly upregulate *Scg2* expression in primary cultured astrocytes at 2 to 8 h (**Fig. 6j**), with increased intracellular calcium and cFos expression (**Supplementary** Fig. 6a, b). SCG2 protein was actually secreted into the medium and its levels were elevated by ATP (**Fig. 6k**, **l**). We further confirmed that extracellular ATP levels were significantly higher in the denervated side of the cervical cord than in the intact side at day 3 to 14 post-stroke (**Fig. 6m**, **n**). Collectively, the results suggest that extracellular ATP increased in the denervated cervical cord stimulates *Scg2* expression and secretion in astrocytes after stroke.

### *Scg2* expression is evoked by neural activity in V2a interneurons and astrocytes

Previous studies showed that neural activity in corticospinal neurons or spinal interneurons promotes axon reorganization and recovery (Lee et al., 2011; Wahl et al., 2017; Van Steenbergen et al., 2023; Yang and Martin, 2023). Rehabilitative training, which elevates neural activity, also enhances reorganization and recovery (Nakagawa et al., 2013; Okabe et al., 2016; Tanaka et al., 2020). We therefore hypothesized that the molecules that promote reorganization should be upregulated during rehabilitation. We used a rotarod rehabilitation to promote axon reorganization (**Fig. 7a**) (Nakagawa et al., 2013). The rehabilitation initiated on day 7 post-stroke increased axon density at day 14 (**Fig. 7a–c**). The rotarod increased the expression of immediate early gene cFos in V2a neurons (**Fig. 7d**), suggesting elevated activity. To explore the possibility that the activated V2a neurons and surrounding astrocytes might increase the expression of molecules for reorganization, we again conducted RiboTag and RNA-seq in *Chx10*- *Cre*;*RiboTag* and *Aldh1l1*-*CreER*;*RiboTag* mice at 2, 4, and 8 h after rotarod exercise on day 14 post-stroke. PCA showed high similarity in gene expression across the groups (**Supplementary** Fig. 7a). After the exercise, we found that a number of transcripts were increased in V2a neurons and astrocytes (**Fig. 7e**, **f**, **Supplementary** Fig. 7b, c). Notably, *Scg2* was found to be upregulated in astrocytes 4 h after the exercise, which was further confirmed by quantitative PCR (**Fig. 7g**, **h**). Furthermore, *Scg2* expression also increased in V2a neurons at 2 to 8 h (**Fig. 7i**, **j**).

**Figure 7.**
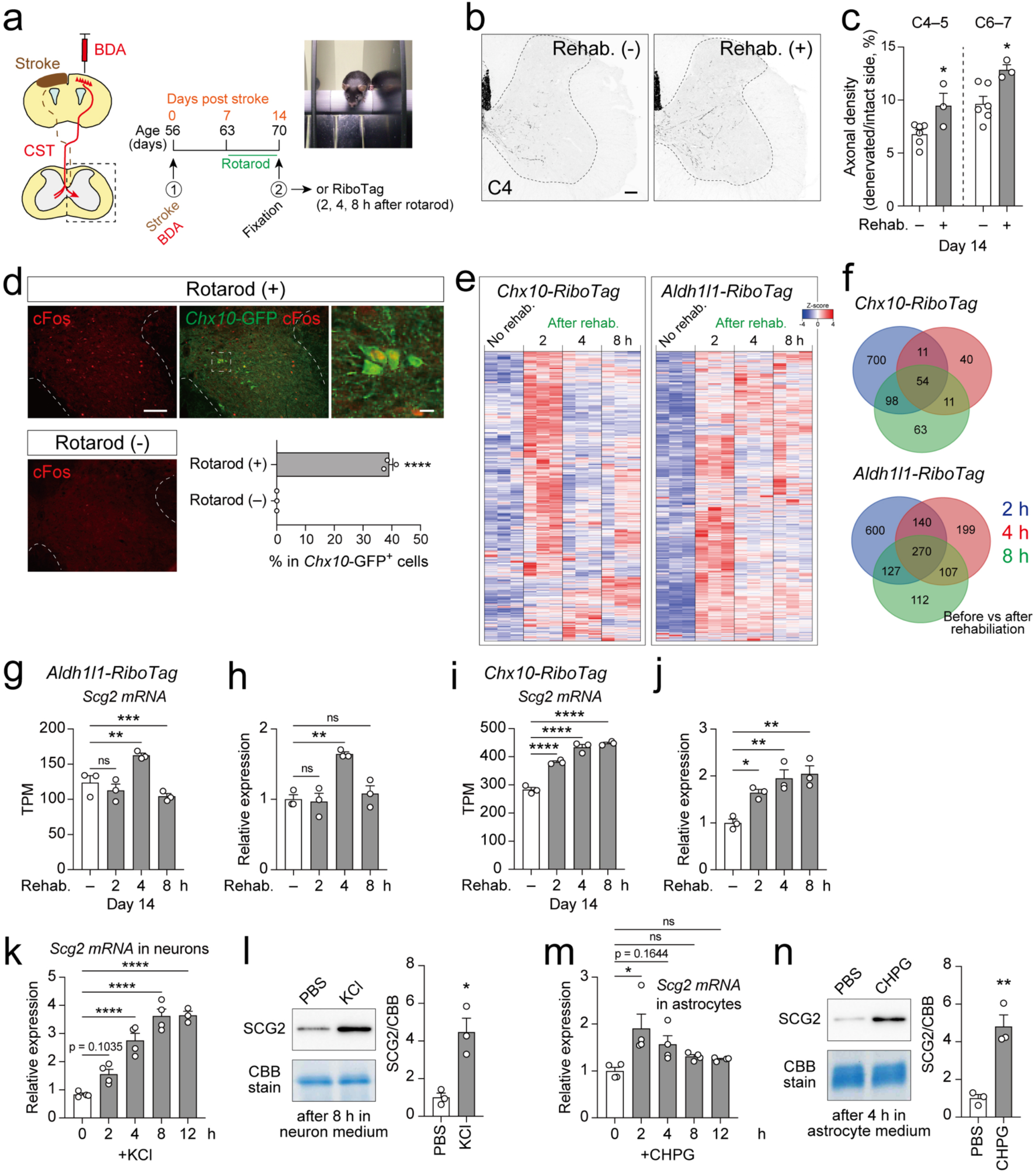
Neuronal activity induces *Scg2* expression in V2a interneurons and astrocytes. **a**, Experimental diagram to assess CST reorganization and RiboTag gene expressions in mice with rehabilitative training by rotarod. Anterograde CST labeling from the contralesional motor cortex (left), experimental timeline (middle), and a photograph of a rotarod (right). **b**, Representative BDA^+^ CST axons in the denervated side at day 14 post-stroke with (right) or without rotarod rehabilitation (left, C4 level). The dotted lines represent the boundary between the gray and white matter. **c**, Quantification of CST axon density in the denervated side by rotarod rehabilitation. N = 6 (no rehabilitation), 3 (rehabilitation); Student’s *t*-test, \**p* < 0.05. **d**, Representative images of cFos expression in GFP*^+^* V2a neurons in *Chx10-Cre;lcl-EGFP* mice with and without rotarod rehabilitation. Bar graph indicates the ratio of cFos^+^ cells in *Chx10*-GFP^+^ neurons. N = 3 mice, Student’s *t*-test, \*\*\*\**p* < 0.0001. **e**, Heatmap showing upregulated DEGs at 2–8 h after rotarod rehabilitation in *Chx10-Cre*;*RiboTag* and *Aldh1l1-CreER*;*RiboTag* mice. N = 3 mice for each group. **f**, Venn diagrams showing the number of upregulated DEGs after rotarod rehabilitation. **g**–**j**, Expression levels of *Scg2* mRNA in astrocytes (**g**,**h**) and V2a neurons (**i**,**j**) after rehabilitation in RNA-seq (**g**,**i**) and quantitative PCR (**h**,**j**). N = 3, multiple testing by Benjamini and Hochberg procedure (**g**,**i**), one-way ANOVA followed by Tukey’s test (**h**, **j**). **k**,**l**, Expression of *Scg2* mRNA (**k**) and secretion of SCG2 protein in conditioned media (**l**) after TTX/KCl treatment in cultured cortical neurons. N = 4, one-way ANOVA followed by Tukey’s test (**k**). Right graph of **l**, quantification of SCG2 protein levels in the medium. N = 3, Student’s *t*-test. **m**,**n**, Expression level of *Scg2* mRNA (**m**) and secretion of SCG2 protein in conditioned media (**n**) after CHPG treatment in cultured astrocytes. N = 4, one-way ANOVA followed by Tukey’s test (**m**). Right graph in **n**, quantification of SCG2 protein levels in the medium. N = 3, Student’s *t*-test. Scale bars, 100 µm (**b**, left in **d**); 10 µm (right in **d**). ns, not significant; \**p* < 0.05, \*\**p* < 0.01, \*\*\**p* < 0.001, \*\*\*\**p* < 0.0001.

Since Scg2 was previously shown to be regulated by neural activity and Fos (Nedivi et al., 1993; Yap et al., 2021), enhanced neuronal activity through rehabilitation might induce *Scg2* expressions in V2a neurons as well as in astrocytes. We examined whether enhanced neuronal activity actually induced *Scg2* expression in neurons. Increasing the activity by the voltage-dependent Na^+^ channel inhibitor TTX followed by KCl administration increased *Scg2* expression in cultured cortical neurons from 2 to 12 h (**Fig. 7k**). Additionally, secretion of SCG2 protein was also elevated (**Fig. 7l**). Enhanced neural activity also increases neurotransmitter release such as glutamates. Astrocytes surrounding synapses respond to glutamates through metabotropic glutamate receptor 5 (mGluR5), which trigger intracellular calcium signals (Bazargani and Attwell, 2016; Dallérac et al., 2018). To assess the response of astrocytes to glutamate signals, we stimulated cultured astrocytes with CHPG, a selective agonist for mGluR5. We found that *Scg2* expression transiently increased 2 h after CHPG treatment, along with SCG2 protein secretion (**Fig. 7m**, **n**). CHPG also increased the intracellular calcium concentration and cFos expression (**Supplementary** Fig. 7d, e), suggesting an induction of signaling pathways similar to those activated by ATP stimulation (**Supplementary** Fig. 6a, b). Collectively, the results suggest that enhanced neural activity promotes the expression and secretion of SCG2 in V2a neurons and astrocytes, which may engage in axon reorganization.

### SCG2 enhances CST rewiring after stroke

Finally, we examined whether SCG2 drives CST axon reorganization in vivo. To overexpress SCG2, we generated an AAV vector expressing HA-tagged SCG2 (AAV-CAG-Scg2-HA) and confirmed that the cultured cells infected with the AAV efficiently secreted SCG2 (**Supplementary** Fig. 8a). The administration of collected supernatant medium promoted phosphorylation of S6 similar to the effect observed with recombinant SCG2 protein in CGNs (**Supplementary** Fig. 8b). We then injected AAV-CAG-Scg2- HA or control AAV-CAG-HA into the denervated side of the cervical cord at day 7 post-stroke and examined if SCG2 overexpression would enhance the reorganization (**Fig. 8a**). We found that the overexpression significantly increased axon density in the denervated side after stroke (**Fig. 8b–d**). The axon density was increased in the medioventral area including laminae V–VIII (**Fig. 8b**, **c**) and lateral and dorsoventral extensions were promoted, especially beyond 400 µm from the midline (**Supplementary** Fig. 8c–f). Under this condition, we also found that S6 phosphorylation levels were elevated in CTIP2^+^ layer V neurons in the sensorimotor cortex ipsilateral to the injection side (**Fig. 8e**, **f**; **Supplementary** Fig. 8g, h).

**Figure 8.**
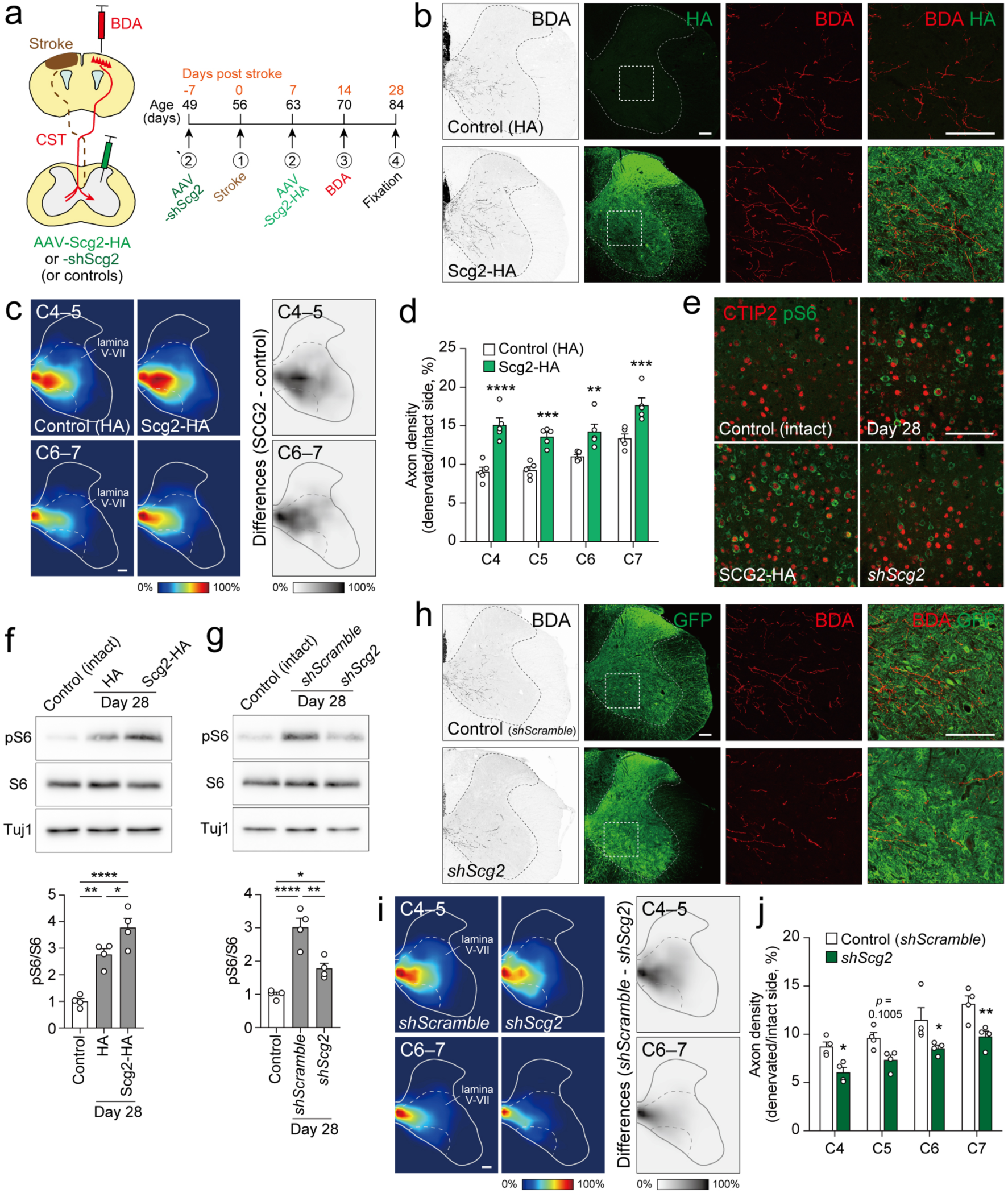
SCG2 promotes rewiring of CST axons after stroke. **a**, Experimental diagram of CST tracing by BDA injection in the motor cortex and SCG2 overexpression or knockdown by AAV injection into the cervical cord. Right, experimental timeline. **b**, Representative images of BDA^+^ CST axons and AAV-derived SCG2-HA expression (green) in the denervated side of the cervical cord at day 28 post-stroke. Right four panels show higher magnification views of the dotted squares in HA staining images. Note that a short HA coding sequence in control AAV did not show detectable levels of HA expression. **c**, Heatmaps showing the axon distribution of CST (left) in control and SCG2 overexpressing mice at day 28 post-stroke (left) and the difference in axon density (subtraction of the controls from the SCG2 overexpressing group, right). **d**, Quantification of CST axon density in the denervated side of control and SCG2 overexpressing cervical cord at day 28 post-stroke. N = 5, two-way ANOVA followed by Sidak’s post hoc test. **e**, Representative confocal images of phospho-S6 (green) and CTIP2^+^ layer V neurons (red) in intact control and stroke mice at day 28 with SCG2 overexpression and knockdown. Enlarged views of each image are in Supplementary Fig. 8h. **f**,**g**, Western blots showing phosphorylation of S6 proteins in the sensorimotor cortex of control and stroke mice with SCG2 overexpression (**f**) and knockdown (**g**). Schematic representation of the experiment is provided in Supplementary Fig. 8g. Bottom, quantification of phospho-S6 expression levels. N = 4, one-way ANOVA followed by Tukey’s test. **h**, Representative images of BDA^+^ CST axons and control shRNA or *Scg2* shRNA-induced cells (GFP^+^, green) in the denervated side of the cervical cord at day 28 post stroke. Right four panels represent higher magnifications of the dotted squares in GFP staining images. **i**, Heatmaps showing the axon distribution of CST in control *shScramble* and *shScg2* induced mice at day 28 post-stroke (left) and the differences in axon density (subtraction of the *shScg2* groups from control *shScramble* group, right). **j**, Quantification of CST axon density in the denervated side of *shScramble* and *shScg2* induced cervical cord at day 28 post-stroke. N = 4, two-way ANOVA followed by Sidak’s post hoc test. Scale bars, 100 µm. \**p* < 0.05, \*\**p* < 0.01, \*\*\**p* < 0.001, \*\*\*\**p* < 0.0001.

We next investigated whether *Scg2* knockdown inhibits the axon reorganization. We generated *Scg2* shRNA-expressing AAV using an miR30-based shRNA system (AAV-CAG/GfaABC1D-GFP-miRE-*shScg2*) (Fellmann et al., 2013). Two of the three *shScg2* constructs exhibited a high knockdown efficiency in Neuro2a cells (**Supplementary** Fig. 8i, j). We selected *shScg2*-#3 for knockdown and infused the AAV into the denervated side of the cervical cord (**Fig. 8a**). *Scg2* knockdown was found to decrease CST axon density at most the cervical levels at day 28 (**Fig. 8h–j**), while the control *shScrambled* did not. Heatmaps and density quantification showed a reduced proportion of laterally and ventrally extending axons (**Fig. 8i**, **Supplementary** Fig. 8k–m). We also found that *Scg2* knockdown inhibited S6 activation in layer V neurons ipsilateral to the injection side (**Fig. 8e**, **g**, **Supplementary** Fig. 8g, h). Taken together, the data revealed that SCG2 is required to induce CST axon rewiring after stroke.

## Discussion

The connectivity of the reorganized circuits for recovery and the molecular mechanisms governing the reorganization are poorly understood in CNS injuries. The present study showed that rewired CST axons reformed functional connections with V2a spinal interneurons and they are essential for restoring skilled motor function after stroke. We further revealed the molecular mechanisms inducing the reorganization, in which degeneration and neural activity in the denervated area upregulate Scg2 in astrocytes and target neurons and it drives regrowth of surviving axons to reorganize the circuit (**Supplementary** Fig. 9).

## Neural substrate for motor recovery

The detail circuit that the CST reformed for recovery after injuries were not well understood. While specific rehabilitation tasks or molecular approaches have succeeded in promoting growth of CST axons, aberrant growth often leads to maladaptive changes or absence in recovery, possibly due to forming a malfunctional connections (Fawcett, 2009; Lu et al., 2012; Tan et al., 2012; Nakagawa et al., 2013; Wahl et al., 2014; Geoffroy et al., 2015; Wang et al., 2015). This implies that it is crucial for CST axons to connect with appropriate target neurons to form functional circuits for recovery. V2a neurons are premotor spinal neurons that control skilled movement as well as locomotion in intact mice (Crone et al., 2008; Azim et al., 2014b; Ueno et al., 2018). Previous studies showed insights that some CST collaterals had anatomical contacts with Chx10^+^ neurons after pyramidotomy or spinal cord injury in rodents and monkeys (Liu et al., 2015; Nakagawa et al., 2019; Van Steenbergen et al., 2023). However, it remained unclear whether they reflect essential and abundant connections and functions in the rewired circuit. By comprehensively studying the locations of interneuron subtypes and employing genetic labeling, synaptic markers, in vivo fiber photometry with optogenetics, and chemogenetic silencing in 3D motor analyses, we provided strict evidence that V2a neurons are key spinal neurons that reconstituted connections with CST axons for motor recovery. The reorganized V2a circuit would complement the lost circuits and functions after stroke. The connection and functional changes observed between the CST and V2a neurons, even in mid-to-later stages, may reflect the ongoing refinement or re-learning process of the behavior by a remnant circuit that was altered after the stroke (Hwang et al., 2019). Recent reports in spinal cord injury also showed the involvement of V2a neurons of the cervical to lumbar cord in the recovery of hindlimb locomotion (Bradley et al., 2019; Kathe et al., 2022; Squair et al., 2023; Van Steenbergen et al., 2023). Propriospinal neurons, which possibly include V2a population, have also been shown to be responsible for dexterous hand movement recovery in CST injuries in primates (Tohyama et al., 2017). These support the importance of this neuronal subtype as a central hub neuron to achieve circuit reorganization and motor recovery in a wide range of CNS injuries.

The experiments of neuronal silencing indicate that V2a neurons are recruited for the recovery of the motor aspect, especially in advancing the forelimb. However, motor control requires coordinated activity of excitatory and inhibitory spinal neurons to regulate extensor and flexor muscle movements with sensory feedbacks (Goulding, 2009; Azim et al., 2014a). Thus, reconnection with other neuronal subtypes would also be involved or required to fully achieve the recovery. Indeed, the success parameters were not fully abolished just by silencing the V2a neurons. The correlation-coefficient analysis suggests the involvement of additional target populations such as Isl1^+^ (dl3), Dbx1^+^ (V0/dl6), and Nkx2.2^+^ (V3) neurons. Further studies are required to better understand the spinal elements as well as other descending and sensory feedback pathways involved in the recovery process. In this context, while the functional connections have mainly been analyzed in acute slices (Ueno et al., 2018), the current photometric method succeeded in recording the responses of specific spinal neurons in vivo. This will allow to analyze the network of other spinal neurons in future studies.

## Molecular mechanisms for axon rewiring

Injuries induce axon denervation in diverse target areas; however, the molecular mechanisms that drive the axon reorganization have long remained unknown (Liu and Chambers, 1958; Raisman, 1969; Tsukahara, 1981). The process includes sequential cellular changes in the denervated area, which may also be induced commonly in different target areas and types of injury (**Supplementary** Fig. 9). Our translated RNA-seq data revealed that the cells in the remote denervated area responded well to the injuries, suggesting that the cells and their expressed molecules engage in circuit repair. Importantly, the identification of target neurons in our analyses allowed us to examine gene expressions in specific neurons involved in the reorganized circuit. While several factors, such as BDNF, CNTF, and GDF10, have been shown to be involved in the axon rewiring (Li et al., 2010; Ueno et al., 2012; Jin et al., 2015; Chang et al., 2022), the central molecular mechanisms triggering this process remained unknown. Since at least 40% of the translational response is independent with transcription levels (Schwanhäusser et al., 2011; Sapkota et al., 2022), the present database of translated mRNA analyses, conducted in a cell type-specific manner, would contain previously unknown candidate genes, which might not be detectable even with the current single-cell RNA-seq technologies.

Our data propose that Scg2 serves as one of the master regulators causing reorganization, especially in response to injury signals and neural activity (**Supplementary** Fig. 9). Elevated extracellular ATP levels by the injury induced expression of Scg2, suggesting ATP as the first trigger of the reorganization. Indeed, damage-associated molecular patterns (DAMPs) released from injured cells, particularly ATP, are implicated in astrocyte and microglial activation (Wang et al., 2004; Davalos et al., 2005; Burda et al., 2016; Bouras et al., 2022). Glial cells in the distal areas possibly respond to signals from injured axons that originate the lesion and secrete molecules that support circuit repair such as for axonal extension and synaptogenesis. The ATP–SCG2 axis may play a common role in the degenerative areas, where circuit remodeling occurs in a wide range of CNS disorders. SCG2 may function to transform the damaged environment from inhibitory to supportive for axon growth, rather than controlling axon guidance or specificity of connections, which should be further studied even in the developmental process. Especially, an increase in intracellular cAMP would shift the axons to a growth state on the inhibitory substrates such as myelin-related proteins as well as to a responding state to neurotrophic factors such as BDNF (Song and Poo, 1999; Cai et al., 2001; Ming et al., 2001). Elevated pS6 levels associated with increased translations would also contribute to axonal growth as observed in *Pten* knockouts (Liu et al., 2010; Nakamura et al., 2021).

In addition to its direct effects on neurons presented in this study, SCG2 regulates the secretory pathway by sorting and packaging growth factors, neuropeptides, and hormones into DCVs (Taupenot et al., 2003; Bartolomucci et al., 2011). Elevated SCG2 increases the number of DCVs and enhances the secretory activity (D’Alessandro et al., 2008; Prada et al., 2011; Lin et al., 2022). Since our results show an increased number of SCG2^+^ puncta in astrocytes, SCG2 might also enhance the secretory pathway after stroke. Notably, BDNF, which induces axon rewiring of the CST (Ueno et al., 2012), is also stored in DCVs in neurons as well as astrocytes for secretion (Verkhratsky et al., 2016; Hoogstraaten et al., 2020; Moro et al., 2020; Han et al., 2021). Our RNA-seq data actually showed elevated *Bdnf* in V2a neurons after rehabilitative training (data not shown) together with *Scg2*. Thus, secretory molecules may function cooperatively in circuit repair in the secretory machinery, which is centrally regulated by Scg2. Comprehensive profiling of the secretome may further elucidate the reparative mechanisms of the circuit following CNS injuries.

In conclusion, the present study demonstrated specific spinal population critical for reorganizing the motor circuit and restoring motor function after stroke. The molecular mechanism governing the reorganization process was further identified. These findings highlight the neural and molecular bases controlling circuit reorganization and recovery and they could be potential therapeutic targets to enhance functional recovery in various types of CNS injury.

## Methods Animals

C57BL/6J (Charles River), *Chx10*-*Cre* (a gift from S. Crone, CCHMC, and K. Sharma, University of Chicago (Azim et al., 2014b)), *Aldh1l1*-*CreER* (Jackson laboratory, #029655), *Cx3cr1*-*CreER* (Jackson laboratory, #020940), *CAG*-*lox*-*CAT*-*lox*-*EGFP* (*lcl*- *EGFP*; a gift from J. Robbins, CCHMC (Nakamura et al., 2006)), and *floxed*-*Rpl22^HA^*mice (*RiboTag*; Jackson laboratory, #011029 (Sanz et al., 2009)) were used. They were housed with light on a 12-h cycle and food and water *ad libitum*. All experimental procedures were performed in accordance with the protocol approved by the Institutional Animal Care and Use Committee of Niigata University.

## Plasmids

The following plasmids were generated for AAV production or transfections in culture: for pAAV-Syn-DIO-jGCaMP7s and pAAV-CAG-jGCaMP7s, jGCaMP7s fragment with BamHI/HindIII site derived from pGP-AAV-Syn-jGCaMP7s-WPRE (Addgene, #104487) was subcloned into pAAV-Syn-DIO-MCS, which had multicloning sites (MCS) replacing mCherry in pAAV-Syn-DIO-mCherry (Addgene, #50459), and into pAAV-CAG-tdTomato (Penn vector core; AV-1-PV3365) in a substitution of tdTomato, respectively. For pAAV-CAG-Scg2-HA and pAAV-CAG-HA, the coding sequence of mouse *Scg2* fused to HA-tag at the C-terminal was amplified from cDNA of mouse spinal cord of 8-weeks old using F1/R1 primers (Supplementary Table 1). HA-tag fragment was prepared by annealing oligonucleotides (oligo-1; Supplementary Table 1). They were inserted into the BamHI and HindIII sites of pENN-AAV-CAG-tdTomato-WPRE-SV40 (Addgene, #105554). For pAAV-CAG-EGFP-shScg2 and pAAV-CAG-EGFP-shScramble, the coding sequences for *shScg2* and *shScramble* were amplified from oligonucleotides (oligo-2, 3, 4, and 5 for *shScg2-*#1, #2, #3 and *shScramble*, respectively) and F2/R2 primers (Supplementary Table 1) and were cloned into the XhoI and EcoRI sites of pAAV-Ptet-RFP-shR-rtTA (Addgene, #35625), which encodes miR-30-based shRNA expression system (shR). The RFP-shR fragment was subcloned into the BamHI and HindIII sites of pENN-AAV-CAG-tdTomato-WPRE-SV40 to generate pAAV-CAG-RFP-shScg2 and pAAV-CAG-RFP-shScramble. The RFP sequence was replaced with EGFP, which was amplified from pAAV-Syn-EGFP (Addgene, #50465) using F3/R3 primers (Supplementary Table 1). For pZac2.1-gfaABC1D-EGFP-shScg2 and pZac2.1-gfaABC1D-EGFP-shScramble, the EGFP-shScg2 and EGFP-shScramble fragments were subcloned into the BamHI/NotI site of pZac2.1-gfaABC1D-cyto- GCaMP6f (Addgene, #52925) in a substitution of GCaMP6f, respectively. For pAAV-Syn-cAMPinG1-NE, cAMPinG1-NE sequence with BamHI and HindIII sites was amplified from pCAG-cAMPinG1-NE (a gift from M. Sakamoto and T. Yokoyama, Kyoto Univ (Yokoyama et al., 2024)) and inserted into pAAV-Syn-ChRmine-mScarlet (Addgene, #130994) in a substitution of ChRmine-mScarlet. For pAAV-Syn-FLEX- oTVBL-mRaspberry-P2A-flag-tTA2S plasmid, oTVB-L (Suzuki et al., 2019), mRaspberry, P2A, and FLAG-tagged tTA2S sequence were inserted into pAAV-Syn-FLEX-MCS. pAAV-TRE-Tight-FLEX-CVSG was constructed by replacing the Syn promoter with the TRE-Tight promoter (Clontech) and inserting G gene of CVS strain rabies virus. pAAV-Syn-ChrimsonR-tdT (#59171) and pAAV-Syn-FLEX-PSAM^4^- GlyR-IRES-EGFP (#119741) were obtained from Addgene.

## Production of AAV vectors

AAV vectors were generated as previously described (Kimura et al., 2023; Nakamura et al., 2023). Briefly, AAVpro 293T cells (632273, TAKARA) were transfected with helper (pHelper; TAKARA), rep/cap (pAAV2/1 and pAAV2/8 for serotype 1 and 8, respectively; Penn vector core), and pAAV plasmids. AAV particles in the cells and supernatants were collected and purified by an iodixanol gradient. The AAVs were then concentrated in phosphate-buffered saline (PBS) containing 0.001% Pluronic F68 using Amicon Ultra 100K filter units (Millipore). The titer of AAV was determined by using AAVpro titration kit (TAKARA) and Thermal Cycler Dice Real Time System III (TAKARA). For AAV2.1-Syn-FLEX-oTVBL-mRaspberry-P2A-flag-tTA2S and AAV2.1-TRE-Tight-FLEX-CVSG, pAAV-RC1 and pAAV-RC2 were used for packaging. AAV particles were purified by affinity chromatography (GE Healthcare) and concentrated to 150 μl by ultrafiltration (Amicon Ultra-4 10K MWCO; Millipore). The titer was determined by quantitative PCR with Taq-Man technology (Life Technologies). The titers of AAV used in this study are described in each section.

## Photothrombotic stroke

Cortical stroke was induced by a photothrombotic method as previously described (Labat-gest and Tomasi, 2013; Ueno et al., 2018; Sato et al., 2021). Male mice at 8 weeks old were placed on a stereotactic frame under anesthesia with isoflurane. The skull was exposed and cleaned, and then a small piece of foil with an opening for light illumination was positioned on the skull over the left sensorimotor cortex (coordinates: mediolateral [ML] 0.5–3.5 mm and anteroposterior [AP] -2.0–3.5 mm from the bregma). Rose Bengal in saline (5 mg/ml, 50 mg/kg body weight, Sigma-Aldrich, 330000) was injected intraperitoneally. Five minutes after the injection, the skull was illuminated with a cold light source (90% output, CL 6000 LED, Zeiss) for 4 min. After the light application, the scalp was sutured and the mice were returned to their home cages.

## Anterograde labeling of CST axons

Anterograde tracing was performed 14 days before fixation as previously described (Ueno et al., 2012, 2018; Sato et al., 2021). Mice were anesthetized with isoflurane and fixed in a stereotactic frame. The scalp was then opened and three small injection holes were made in the corresponding sites of injection with a 27G needle (coordinates; ML 1.0, AP 0.0 mm; ML 1.0, AP 0.5 mm; ML 1.5, AP 0.5 mm; all at a depth of 0.5 mm from the cortical surface). Biotinylated dextran amine (BDA; 10,000 MW; 10% weight/volume in PBS; D1956, Invitrogen) was injected into the target sites of the right cortical hemisphere (contralesional side) using a glass capillary attached to a Hamilton syringe (0.6 µl per site). After the injections, the scalp was sutured and the mice were returned to their home cages.

## Monosynaptic retrograde tracing with rabies virus

*Chx10*-*Cre* mice were anesthetized with isoflurane and fixed in a stereotactic frame. AAV2.1-Syn-FLEX-oTVBL-mRaspberry-P2A-flag-tTA2S and AAV2.1-TRE-Tight- FLEX-CVSG (2.0 x 10^11^ GC/ml in each; 0.4 µl/site; 0.5 mm in depth) were injected into the right cervical cord at C4 and C6 levels at 9 weeks of age. Fourteen days after the AAV injection, EnvB-coated G-deleted rabies virus CVS strain expressing AcGFP (EnvB- CVSΔG-AcGFP; 2.5 x 10^8^ infectious units/ml; 0.2 µl/site) was injected into the same sites of the cervical cord. After the injection, the back skin was sutured and the mice were returned to their home cages. Mice were perfused 7 days after the last injection and histological analysis was performed as described below.

## Histological examination

The animals were deeply anesthetized and transcardially perfused with 4% paraformaldehyde (PFA) in 0.1 M phosphate buffer (PB). The brain and cervical cord were then dissected and post-fixed in the same fixative overnight at 4°C. The samples were cryoprotected in 30% sucrose in PBS and then embedded in an optimal cutting temperature (O.C.T.) compound (Sakura Finetek). The brain and cervical cord were cryo- sectioned serially in 20-µm or 50-µm thickness (for PKCγ staining). For BDA staining, cervical cord sections were incubated with 0.3% Triton X-100 in PBS for 4 h, followed by Alexa Fluor 568-conjugated streptavidin (1:400, S11226, Invitrogen) in 0.1% Tween 20 in PBS for 2 h at room temperature (RT) (Ueno et al., 2012, 2018; Sato et al., 2021). For immunohistochemistry, the sections were incubated for 2 h with 1% bovine serum albumin (BSA) in 0.3% Triton X-100/PBS for blocking, followed by overnight incubation of primary antibodies at 4°C. Rabbit anti-PKCγ (1:500; sc-211, Santa Cruz Biotechnology), rabbit anti-GFP (1:1000; A11122, Invitrogen), rat anti-GFP (1:1000, 04404-84, Nacalai Tesque), goat anti-mCherry (1:2000; AB0040, Sicgen), rat anti-HA (1:500; 11867423001, Roche), rabbit anti-cFos (1:1000; 2250, Cell Signaling), rabbit anti-SCG2 (1:1000; MSRF105440, Nittobo Medical), rat anti-CTIP2 (1:500; ab18465, Abcam), and rabbit anti-pS6 (1:1000; 2217, Cell Signaling) antibodies were used. After washing with 0.1% Triton X-100/PBS, the sections were incubated with corresponding secondary antibodies, Alexa Fluor 488, 568, 647 donkey anti-rabbit, rat, goat or guinea pig IgG antibody (1:1000; Invitrogen or Jackson ImmunoResearch) for 2 h at RT. Images were acquired by using a fluorescence microscope (BX51, Olympus) or confocal microscope (FV3000, Olympus).

For the analyses of anatomical connection of CST axons and Chx10^+^ V2a interneurons, post-fixed cervical cords were immersed in PBS and embedded in low-melting agarose. Serial sections at 80 μm-thick were prepared with a vibratome (VT- 1000S, Leica). Floating sections were blocked with 1% BSA/0.3% Triton X-100 in PBS for 2 h and were then incubated with guinea pig anti-VGLUT1 (1:10000; AB5905, Millipore) and rabbit anti-GFP (1:1000; A11122, Invitrogen) antibodies and Alexa Fluor 568-conjugated streptavidin (1:400) in blocking buffer for 3 overnights at 4°C. After washing with 0.1% Triton X-100 in PBS, the sections were incubated with the corresponding secondary antibodies, Alexa Fluor 488, 647 donkey anti-rabbit, and guinea pig IgG antibody (1:1000; Invitrogen or Jackson ImmunoResearch) for 3 overnights at 4°C. Images were acquired by using a confocal microscope (FV3000, Olympus).

For counting the number of anatomical contacts of BDA^+^ CST fibers onto GFP^+^ V2a interneurons, confocal images were reconstructed by IMARIS software (version 9.5.1, Bitplane). The surface of BDA^+^ axons was first constructed with a smooth parameter of 0.5 µm and VGLUT1^+^ signals within the surface were extracted as spots with a diameter of 3.0 µm. Next, the surface of BDA^+^ axons was reconstructed with a smooth parameter of 1.0 µm and GFP^+^ signals within the surface were detected as spots with a diameter of 3.0 µm. The spots within 3.0 µm from the BDA^+^ surface were further extracted. Finally, VGLUT1^+^/BDA^+^ and BDA^+^/GFP^+^ spots within the distance of 2.75 µm were extracted as anatomical contacts. The number of contacts was measured in 8–20 images per mouse. For analysis of monosynaptic retrograde tracing with the rabies virus, brain and cervical cord sections were prepared in a 50 µm-thickness every 100 µm. The number of AcGFP^+^ input neurons in the motor cortex labeled by anti-GFP antibody was counted in 30 sections covering the area from 2500 µm anterior to 500 µm posterior to the bregma. The number was normalized by the number of AcGFP and mRaspberry3 double-positive starter cells in the cervical cord labeled with anti-GFP and mCherry antibodies, which were counted in 10 sections at each cervical level (C4–7).

## Quantification of the number of crossing axons and axon density

To quantify CST axon rewiring, we counted the number of crossing fibers and measured the axonal density in 10 sections at each cervical cord level, as described previously (Sato et al., 2021). To normalize differences in BDA-labeling among the animals, each value was divided by the mean number of BDA-labeled CST fibers counted in two serial sections of the dorsal column at C4. The axon density in each vertical bin or dorsal and ventral laminae was measured from defined regions of interests (ROIs) in each image by Fiji/ImageJ software (NIH). ROIs for the dorsal and ventral laminae were manually defined according to the mouse spinal cord atlas (Watson et al., 2009). The threshold was adjusted to maintain BDA-positive signals and the images were binarized. The number of pixels corresponding to the BDA-positive signals was measured at each cervical level.

## Heatmap for the axonal density and correlation analysis

Heatmaps of the axonal density were generated as previously described (Sato et al., 2021). The images of the cervical hemicords were binarized and the XY coordinates of pixels corresponding to BDA-labeled signals were measured by Fiji/ImageJ. The coordinates of the central canal positions were determined as the reference points of the coordinates. The adjusted XY coordinates of each image were combined at C4–5 or C6–7. A contour heatmap for axonal density was then generated by MATLAB software (MathWorks).

Heatmaps of interneuron distributions were generated from the data of our previous study (Ueno et al., 2018). The locations of the cell bodies of GFP-labeled interneurons at C4–5 and C6–7 were plotted by Fiji/ImageJ and the XY coordinates were adjusted by the reference points of the central canal. The contour heatmaps of their distributions were generated as described above.

For correlation coefficient analysis, two-dimensional histograms with 24 and 32 bins in x and y-axis, respectively (each 50 µm wide), were generated from the XY coordinate data of BDA^+^ signals and the interneuron locations by MATLAB software. To normalize the two-dimensional histogram data, the values in each bin were divided by the maximum value of all bins. The data were then converted into one-dimensional data in the order of x1y1 to x24y32 and Pearson correlation coefficient was calculated by MATLAB software.

## Recording calcium activity and optogenetic stimulation

AAV was injected into the cervical cord or the cerebral cortex as described previously (Ueno et al., 2018). Mice were anesthetized with isoflurane and fixed in a stereotactic frame. AAV8-Syn-DIO-jGCaMP7s (4.2 x 10^12^ GC/ml; 0.4 µl/site; 0.5 mm in depth) were injected into the right cervical cord at C4 and C6 levels of *Chx10*-*Cre* mice at P14. Then, two weeks before recording of calcium activity, AAV1-Syn-ChrimsonR-tdTomato (5.0 x 10^12^ GC/ml; 0.6 µl; ML 1.0, AP 0.5 mm, depth 0.5 mm) was injected into the right (contralesional) cortex (**Supplementary** Fig. 3a). After the injection, the back skin or the scalp was sutured and the mice were returned to their home cages.

For fiber photometry, mice of controls or at day 28 and 56 after stroke were anesthetized with ketamine/xylazine (100 mg/kg and 10 mg/kg body weight, respectively, i.p.) and placed on a stereotactic frame. A laminectomy was performed at C5 and C7, and the cervical cord at C3 and T2 levels was stabilized with a spinal cord clamp (STS-A, Narishige) to avoid noise from respiration and heartbeats. An optical fiber (200-µm core, NA0.57, Doric lenses) tethered to a low-autofluorescence patch cord (200-µm core, NA0.57, Doric lenses) was held by a stereotaxic arm and attached to the surface of C5 or C7 cervical cord of the denervated side (ML 0.5 mm). Continuous 470 nm excitation lights (M470F3, Thorlabs) were delivered and the emitted fluorescence GCaMP7s signals were collected through the integrated Fluorescence Mini Cube (iFMC5-G2_IE(400- 410)_E(460-490)_F(500-540)_O(580-680)_S, Doric lenses) and recorded by PowerLab (1 kHz sampling rate for 10 sec with a 3-sec pre-stimulus period and a 50-ms photostimulation period, AD instruments). For optical stimulation, a single pulse of 590 nm light (50 ms, 10 mW/mm^2^) was delivered to the recording site through the same fiber by an LED light source (M590F3, Thorlabs). The light exposure and recording were controlled by a stimulus generator (STG4002, Multi Channel Systems). Fifty trials were recorded at a 1-min interval for each mouse.

For control experiments, the cervical cords of *Chx10*-*Cre*;*lcl*-*EGFP* mice injected with AAV1-Syn-ChrimsonR-tdTomato into the right sensorimotor cortex were recorded 14 days after the injection (**Supplementary** Fig. 3b–d). Fluorescence signal changes were not observed, indicating that the setup did not detect noise signals derived from respiration, heartbeats, or other unexpected movements of the animals.

## Data analysis for fiber photometry

Calcium signals were analyzed by MATLAB software with custom-written scripts, as previously described with minor modifications (Sato et al., 2020). The raw data was processed with a moving average filter (10 frames average, 10 ms) and the fluorescence change values (ΔF/F) were calculated by (F_t_ – F_0_)/F_0_ (F_t_, the signal intensity at the recorded time point; F_0_, the baseline intensity averaged in a prestimulation period [0.5–2.5 s before the onset of the stimulation]). The peak amplitude was defined as the maximum value after the onset of optogenetic stimulation. The peak amplitudes of 50 trials in each animal were averaged for quantification. A trial-by-trial heatmap of ΔF/F was generated by MATLAB software.

## Single-pellet reaching test

The single-pellet reaching test was conducted as previously reported (Ueno et al., 2018) with some modifications. Five-week-old male mice were food-restricted to maintain 90% of their free feeding weight before training. The training chamber was made with clear Plexiglas with 0.5 cm wide slits on the left, center, and right sides of the front wall. Millet seeds were placed in front of the slit to allow them to reach, grasp, and retrieve through the slit. A five-day training period was determined by pilot studies, in which success rates reached a plateau in most of the mice. During the initial 3 days of training, seeds were placed in front of the center slit and the mice were allowed approximately 50 attempts per day to learn the test. The right forelimb-preferred mice were selected and 20 reaches per day for the seeds in front of the right slit were recorded for two additional days. The mice that did not learn to reach were omitted for further testing. At 7 weeks of age, reaching tests were conducted again for 3 days. The success rates of the tests were then assessed. When the mouse successfully retrieved the seed and placed it into its mouth, the attempt was considered a success. The quality of each reach attempt was further assessed using a 7-point scale criteria “reaching score”: 6, retrieve the pellet; 5, grasp or touch the pellet; 4, reach beyond the slit; 3, reach into the slit; 2, raise to the slit; 1, raise the forelimb; 0, do not move. The average score of 20 attempts was calculated.

## Chemogenetic inhibition of Chx10^+^ neurons

AAV8-Syn-FLEX-PSAM^4^-GlyR-IRES-EGFP (4.6–6.7 × 10^12^ vg/ml) was injected into C4 and C6 levels of the right spinal cord (0.4 μl / location, 0.4 mm lateral, 0.5 mm depth, right side) of *Chx10*-*Cre* mice at P14. Training of single-pellet reaching test was started at 5 weeks of age and only right-handed mice were used for the experiment. At 7 weeks, the reaching test was repeated for 3 days. On the last day pre-stroke and day 28 and 56 post-stroke, the tests were performed before and 30 min after an injection of uPSEM817 (0.3 mg/kg body weight, i.p.; 6866, Tocris).

## Kinematic analyses of reaching behavior

Kinematic features of forelimb movement during reaching behaviors were assessed with high-speed cameras and a KinemaTracer system (Kissei Comtec) (Ueno and Yamashita, 2011; Ueno et al., 2018; Nakamura et al., 2023). During the reaching test, 8–12 videos of 11-s duration were acquired with two high-speed digital cameras (200 fps, Grasshopper, Point Grey) placed diagonally behind and in front of the slit. Recordings were acquired by the KinemaTracer software, and the movements of the back of the paw (base of the middle finger) were automatically traced by DeepLabCut (Mathis et al., 2018). The KinemaTracer software was modified to import the coordinate data generated by DeepLabCut. Before each session, the precise coordinates were calibrated by recording a cube of a known size (5 × 10 × 10 cm [x × y × z]). The far distal positions of the reaching paw at the time that grasping was initiated were plotted and the relative positions to the pellet were analyzed on the x-y axis. Their spatial probabilities were represented in heatmaps. Since frame-by-frame analyses of motion components revealed abnormalities in the advancement step by inhibition of Chx10^+^ neurons (Ueno et al., 2018), parameters of the average values of acceleration (cm/s^2^) between the time points of minimum and maximum velocities during the period when the forelimb was raised and moved towards the seed (the advancement step) were obtained and compared.

## Tamoxifen injection

To activate CreER recombinase, tamoxifen (T5648, Sigma-Aldrich) was dissolved in corn oil (20 mg/ml) and administrated by intraperitoneal injection (0.1 mg/g body weight, i.p.) at 6 weeks of age. The injection was done once for *Aldl1l1*-*CreER*;*lcl*-*EGFP* mice to sparsely label astrocytes and for two consecutive days for *Aldl1l1*-*CreER*;*RiboTag* and *Cx3cr1-CreER*;*RiboTag* mice, which induced HA expression in most of the astrocytes and microglia, respectively.

## Extraction of cell-type specific mRNA from *RiboTag* mice

*Chx10*-*Cre*, *Aldl1l1*-*CreER*, and *Cx3cr1*-*CreER*;*RiboTag* mice were deeply anesthetized with pentobarbital at days 3, 7, 14, or 28 after stroke. The cervical cord between C4 and C7 was quickly dissected and divided into intact and denervated sides along the midline. The dorsal column containing activated astrocytes and microglia was further removed in *Aldl1l1*-*CreER*;*RiboTag* and *Cx3cr1*-*CreER*;*RiboTag* mice. Dissected tissues were immediately frozen and stored at -80 °C until RNA extraction.

mRNA extractions from V2a interneurons, astrocytes, and microglia were performed as previously described (Sanz et al., 2009, 2019). In brief, the collected tissue was homogenized in HB-S buffer (50 mM Tris-HCL [pH 7.4], 100 mM KCl, 12 mM MgCl_2_, 1% NP-40, 1 mM DTT, 1 mg/mL heparin, 100 µg/mL cycloheximide, 200 U/mL RNasin [Invitorgen], protease inhibitor cocktail [Sigma-Aldrich]) on ice and then centrifuged at 10,000 g for 10 min at 4°C. A small portion (5%) of the supernatant was kept as an input sample, which would contain mRNAs of all the cell types. The remaining supernatant was mixed with 5 µl of rabbit anti-HA antibody (ab9110, Abcam) and incubated for 4 h at 4°C with gentle mixing. Dynabeads protein G (Invitrogen) were then added and the mixture was incubated overnight. To precipitate the RNA/antibody/beads complex, the sample tube was placed on a magnetic stand (DynaMag-2, Invitrogen) until complete precipitation and the supernatant was removed. The non-immunoprecipitated (non-IP) sample was also recovered for a purity check. After washing three times with a high salt buffer (50 mM Tris-HCL [pH 7.4], 300 mM KCl, 12 mM MgCl_2_, 1% NP-40, 0.5 mM DTT, and 100 µg/mL cycloheximide), cell-type-enriched mRNAs from the immunoprecipitation (IP) samples were extracted using the RNeasy Plus Micro kit (Qiagen). The quantity and quality of RNA were assessed by TapeStation with a High Sensitivity RNA ScreenTape (Agilent Technologies). RNAs (10 ng/sample) with RNA integrity number (RIN) values greater than 8.0 were used to prepare libraries for RNA-seq.

## RNA sequencing and analysis

cDNA libraries were prepared using the SMART-seq v4 Ultra Low Input RNA Kit (TAKARA) and Nextera XT DNA Library Preparation Kit (Illumina) according to manufacturer’s instructions. Sequencing was then performed on Illumina NextSeq500 (2 × 75 bp read length). Raw sequence reads were trimmed and their quality was assessed by Trimmomatic (version 0.39; (Bolger et al., 2014)) and FastQC (version 0.11.9) programs, respectively. Trimmed reads were then aligned to the mouse genome (GRCm39/m32) by the STAR aligner (version 2.7.9a; (Dobin et al., 2013)) with default parameters. Approximately 85–90% of the reads were mapped uniquely to the mouse reference genome and used for subsequent analyses. Read counts and transcript per million (TPM) values were calculated using RSEM program (version 1.3.1; (Li and Dewey, 2011)) at default setting. PCA values were computed by the prcomp function in R software and plotted by MATLAB software. To identify differentially expressed genes (DEGs), DESeq2 program (version 1.32.0; (Love et al., 2014)) was used with a false discovery rate (FDR) threshold of < 0.1 and a 1.5-fold change in expression levels (log_2_ fold change > 0.585 or < -0.585). Gene lists of annotation for secretory proteins were obtained by Metascape program (Zhou et al., 2019). Heatmaps displaying DEGs and Venn diagrams showing the number of DEGs were generated by R software.

## Reverse transcription and quantitative PCR

Total RNA was extracted from cultured cells by RNeasy Plus Universal Kit (Qiagen). The RNAs or the ones extracted by the RiboTag method were then reverse transcribed for first-strand cDNA synthesis by the SuperScript III First-Strand Synthesis Kit (Invitrogen) or PrimeScript RT Master Mix (TAKARA). Real-time PCR was performed with TB green fast qPCR Mix (TAKARA) and the primers (Supplementary Table 1) in the Thermal Cycler Dice Real Time System III (TAKARA). The expression level of each mRNA was normalized with that of *Gapdh*.

## Extracellular ATP measurement

Extracellular ATP concentration was measured as previously described (Cao et al., 2013; Masuda et al., 2016) with minor modification. The cervical cord segments (C4–7) were dissected under deep anesthesia with pentobarbital and 500 µm coronal slices were prepared with a vibratome. The slices were then divided into intact and denervated sides at the midline and incubated in ice-cold oxygenated artificial CSF (aCSF) (125 mM NaCl, 2.5 mM KCl, 2.0 mM CaCl_2_, 2.0 mM MgSO_4_, 1.25 mM NaH_2_PO_4_, 26 mM NaHCO_3_, and 10 mM glucose) for 15 min. The aCSF was then collected and ATP levels were assessed by CellTiter-Glo Luminescent Cell Viability Assay (Promega) and a microplate reader (FilterMax F5, Molecular Devices). A standard ATP solution (01072-11, Nacalai Tesque) was used to generate a calibration curve. To normalize the ATP levels among the samples, protein concentration in the aCSF was measured by the BCA assay kit (Thermo Fisher).

## Cell culture

The cells were cultured at 37°C with 5% CO_2_. Cortical neurons and CGNs were prepared as previously described (Ueno et al., 2013; Nakamura et al., 2021) with slight modifications. Briefly, the cerebral cortex and cerebellum were removed from E15 embryo and P7 mice, respectively. The pia mater was peeled off and the tissues were dissociated with a reagent containing 0.25% trypsin and 0.1% DNase I in PBS for 15 min at 37°C. The cells were triturated gently with a pipette to obtain a single cell suspension. The cells were then passed through a 40-μm filter, centrifuged, and resuspended in

DMEM/F12 (Gibco) with 2% B-27 supplement (Gibco) and penicillin/streptomycin (P/S; Invitrogen).

For the neurite outgrowth assay, CGNs were plated on 35-mm dishes at a density of 2 × 10^5^ cells/dish. For preparation of cell lysates, the cells were plated on poly-L-lysine (PLL; 100 µg/ml, 0413, ScienCell Research Laboratories,)-coated 35-mm dishes at a density of 1 × 10^6^ cells/dish. After 2 days of culture, the cells were treated with SCG2 (100 ng/ml; LS-G51092, LifeSpan Biosciences) or conditioned medium derived from Neuro2a cells. After 30 min of incubation, CGNs were lysed and subjected to immunoblotting. For some experiments, SQ22536 (100 µM; Q0105, Tokyo Chemical Industry) was added 1 h before SCG2 treatment. For cAMP imaging, CGNs were transfected with pAAV-Syn-cAMPinG1-NE using Lipofectamine 3000 (Invitrogen) at 2 h after plating. Half of the medium was replaced at 2 days after transfection. The imaging was performed on day 4 after transfection as described below.

To enhance neuronal activity in cortical neurons, the cells were plated on PLL-coated 35-mm dishes at a density of 3 × 10^6^ cells/dish and then TTX (1 µM; 32775-51, Nacalai Tesque) and KCl (55mM) were added on day 4 and 5 after plating, respectively. Cells were extracted at 0–12 h after KCl treatment and the total RNA and conditioned media were collected for immunoblotting.

For astrocytes, the cerebral cortex was removed from P1 mice, and the pia mater was peeled off. Tissues were trypsinized with dissociation reagent for 10 min at 37°C and then triturated gently with a pipette. The cells were passed through a 70-μm filter, centrifuged, and resuspended in the culture medium (DMEM [Wako] containing 10% fetal bovine serum [FBS, Biowest] and P/S). The cells were then plated on 100-mm dishes and cultured until reaching confluent (7–10 days), with changing the medium every 3 days. To remove microglia and oligodendrocytes, the dishes were shaken at 200 rpm for 2 h. Astrocytes were then harvested and replated on 35-mm dishes at a density of 2 × 10^5^ cells/dish. After an overnight culture, the medium was replaced with a flesh medium. The cells were then treated with ATP (100 µM; 01072-11, Nacalai Tesque) or CHPG (100 µM; 1049, Tocris) and incubated for 0–12 h for extraction of mRNA and collection of conditioned media. For cFos staining and calcium imaging, astrocytes were plated on 35-mm dishes at a density of 5 × 10^4^ cells/dish. The cells were then treated with ATP (100 µM) or CHPG (100 µM) and incubated for 1.5 h, followed by fixation with 4% PFA/PB for 30 min. For labeling jGCaMP7s, the cells were transfected with pAAV-CAG-jGCaMP7s with Lipofectamine 3000 (Invitrogen).

Neuro2a cells were maintained in the cultured medium. For conditioned media, the cells were infected with 2 µl of AAV8-CAG-SCG2-HA or AAV8-CAG-HA (2.0 x 10^12^ GC/ml each) in a 35-mm dish. Two days after infection, the medium was replaced with differentiation medium (0.5% FBS and P/S in DMEM), and the medium was further changed to serum-free DMEM/F12 at 7 days post-infection. After 8 h, the serum-free conditioned medium was collected and added to CGNs. To evaluate the knockdown efficiency of *Scg2*, the cells were transfected with pAAV-CAG-EGFP-shScg2 (#1–3) and pAAV-CAG-EGFP-shScramble using polyethylenimine (23966, Polysciences) in 35- mm dishes. They were lysed 72 h after the transfection and then subjected to immunoblotting.

## Neurite outgrowth assay

The neurite outgrowth assay of CGNs was performed as previously described (Nakamura et al., 2021) with slight modifications. The dishes were precoated overnight with PLL and Nogo-A-Fc (3 µl spot, 100 ng/µl; 3728-NG, R&D systems) or CSPG (5 µl spot, 200 ng/ml; CC117, Millipore). CGNs were cultured on the dishes for 2 days. SCG2 (100 ng/ml) was added 4 h after plating. For some experiments, SQ22536 (100 µM) was added 1 h before the SCG2 treatment. CGNs were fixed with 4% PFA/PB for 30 min and subjected to immunocytochemistry as described below. Images were acquired randomly by a fluorescence microscope (BX51, Olympus). Neurite lengths were measured by Fiji/ImageJ software.

## Immunocytochemistry

After fixation with 4% PFA/PB for 30 min, the samples were incubated for 2 h with 1% BSA in 0.1% Triton X-100/PBS for blocking, followed by overnight incubation of primary antibodies at 4°C. Mouse anti-TuJ1 (1:3000; MMS-435P, Covance), chick anti- TuJ1 (1:500; AB9345, Millipore), rabbit anti-cFos (1:3000; 2250, Cell Signaling) and mouse anti-GFAP (1:3000; G3893, Sigma-Aldrich) were used. After washing with 0.1% Triton X-100/PBS, the samples were incubated with the corresponding secondary antibodies, Alexa Fluor 488, 568 donkey anti-mouse or rabbit IgG (1:2000) and Alexa Fluor 568 donkey anti-chick IgY antibody (1:1000; Invitrogen) for 2 h at RT. Images were acquired by a fluorescence microscope (BX51, Olympus).

## Immunoblotting

For the motor cortex, control and stroke mice at day 28 were deeply anesthetized with pentobarbital. Whole brains were quickly dissected and sliced coronally by a brain slicer (MK-MC-01, Muromachi). The contralesional motor cortex was then microdissected. The tissues or the cells were lysed in RIPA buffer (50 mM Tris-HCl [pH 8.0], 150 mM NaCl, 1% NP-40, 0.5% sodium deoxycholate, and 0.1% SDS) containing Protease inhibitors (Roche) and Phosphatase inhibitors (Nacalai Tesque). To analyze secreted proteins in the medium, the supernatants were collected, incubated with 9.09% (v/v) trichloroacetic acid (Nacalai Tesque) on ice for 30 min, and then centrifugated at 15,000 g for 30 min at 4°C to precipitate the proteins. After washing with ice-cold acetone, the precipitates were dissolved in urea buffer (8 M urea, 50 mM Tris-HCl [pH 8.0], and 150 mM NaCl). Immunoblotting was performed by incubation with the following primary antibodies: rabbit anti-pS6 (1:3000; 2217, Cell Signaling), rabbit anti-S6 (1:2000; 2217, Cell Signaling), rabbit anti-pERK (1:2000; 9101, Cell Signaling), rabbit anti-ERK (1:2000; 4695, Cell Signaling), mouse anti-TuJ1 (1:10000; MMS-435P, Covance), rabbit anti-SCG2 antibodies (1:3000; MSRF105440, Nittobo Medical) and rat anti-HA (1:5000; 11867423001, Roche). For secondary antibodies, HRP-linked anti-mouse, rabbit or rat IgG antibodies (1:5000, Cell Signaling) were used. The bands were visualized by Immobilon Forte Western HRP Substrate (Millipore) and LAS3000 (FujiFilm). For CBB staining, Coomassie Brilliant Blue G-250 (C3488, Tokyo Chemical Industry) was used. The intensity of target band was quantified by Fiji/ImageJ.

## cAMP and calcium imaging in vitro

Before imaging, the medium was changed to phenol red-free DMEM/F12 and the cells were incubated at RT for 30 min. cAMPinG1-NE or jGCaMP7s signals were recorded through a water immersion 10× objective lens (N.A. 0.30, UMPLFLN10XW, Olympus) on an upright microscope (BX50, Olympus) equipped with an sCMOS camera (ORCA-Fusion, Hamamatsu Photonics). Images for cAMPinG1-NE and jGCaMP7s were collected at 0.2 Hz (2304 × 2304 pixels) for 6.5 min and 260 Hz (1152 × 1152 pixels) for 3 min, respectively. Excitation light was provided using a 470-nm light-emitting diode module (M470L5, Thorlabs). A standard green fluorescent protein filter set (Olympus) was used to detect the signals. During the imaging, SCG2 (100 ng/ml) was added to CGNs, and ATP or CHPG (100 µM each) was added to astrocytes. The ratio images were generated by Fiji/ImageJ.

## Rotarod rehabilitation

Rehabilitative exercise was performed with a rotarod apparatus with a 30-mm diameter rod (RRAC-3022, O’HARA) as described previously (Nakagawa et al., 2013) with minor modification. Mice were trained in a program where it accelerated from 4 to 40 rpm for 5 min, three times a day, for 3 days before stroke. To analyze axonal density, a rotarod exercise was performed daily for 30 min from day 7 to 14 post-stroke. A constant speed of 10 rpm was applied on the first day and the speed was then gradually increased with one rpm every day. For cFos staining, *Chx10*-*Cre*;*lcl*-*EGFP* mice were run at a constant speed of 20 rpm for 30 min and subjected to fixation 2 h after the exercise as described above. For cFos analysis, cFos^+^ cells in *Chx10*-EGFP^+^ cells were counted in five images at each cervical level (C4–7) for each mouse and the ratio was calculated. For extraction of RNA, *Chx10*-*Cre* or *Aldl1l1*-*CreER*;*RiboTag* mice were trained at a constant speed of 25 rpm for 1 h at day 14 after stroke and then subjected to cervical cord dissection at 2, 4, and 8 h after the exercise as described above.

## Overexpression and knockdown experiments in vivo

BDA and AAV were injected into the cerebral cortex or the cervical cord as described above. For overexpression of SCG2, AAV8-CAG-SCG2-HA or AAV8-CAG-HA (2 x 10^12^ GC/ml in each; 0.4 µl/site; 0.5 mm in depth) were injected into the right cervical cord at C4 and C6 levels at day 7 post-stroke. For knockdown, the 1:1 mixture of AAV8- CAG-GFP-miRE-*shScg2*(#3) and AAV8-GfaABC1D-GFP-miRE-*shScg2*(#3) or AAV8-CAG-GFP-miRE-*shScrambled* and AAV8-GfaABC1D-GFP-miRE-*shScrambled* (4 x 10^12^ GC/ml each; 0.4 µl/site; 0.5 mm in depth) were injected into the right cervical cord at C4 and C6 levels at day 7 before stroke. Mice were perfused at day 28 post-stroke and subjected to analysis of axon rewiring, histology, and immunoblotting as described above.

## Statistical analysis

Statistical analyses were performed with GraphPad Prism 10 software (GraphPad Software). Statistical differences were calculated by a two-tailed unpaired Student’s *t*- test, one-way ANOVA followed by Tukey’s post hoc test and two-way ANOVA with Sidak’s post hoc test. All experiments were conducted with at least three biological replicates. All data are represented as mean ± SEM and significant differences were considered as p < 0.05. Sample sizes are described in the figure legends with statistical methods.

## Acknowledgements

We would like to thank T. Yamashita (Osaka Univ), A. Kakita, K. Shibuki, M. Abe, K. Sakimura, and O. Onodera (Niigata Univ) for supporting materials; S.A. Crone (CCHMC) for providing a mouse line; K. Oda and T. Sasaoka (Niigata Univ) for mouse rederivation; H. Igarashi (Niigata Univ) for kinematic analyses; T. Ikeuchi, N. Hara, S. Miyashita and S. Okuda for RNA-seq analyses (Niigata Univ); M. Sakamoto and T.

Yokoyama (Kyoto Univ) for providing plasmids; K. Ikarashi and R. Furusawa for helping the experiments. This work was supported by JSPS KAKENHI 19K23773 and 21K07290 and Nakatomi Foundation (TS); JSPS KAKENHI 22H04922 and 22H05157 (KI); AMED-CREST (JP21gm1210005), Moonshot Research (JP21zf0127004), JSPS KAKENHI 17H04985, 17H05556, 17K19443, 21H02590, 21H05683, 23H04222, 24K02129, JP 22H04922 (AdAMS), Kato Memorial Bioscience Foundation, Grant-in-Aid from Tokyo Biochemical Research Foundation, Narishige Neuroscience Research Foundation, Ube Industries Foundation, Takeda Science Foundation, and Japan Heart Foundation Research Grant (MU).

---

**Supplementary Figure 1.**
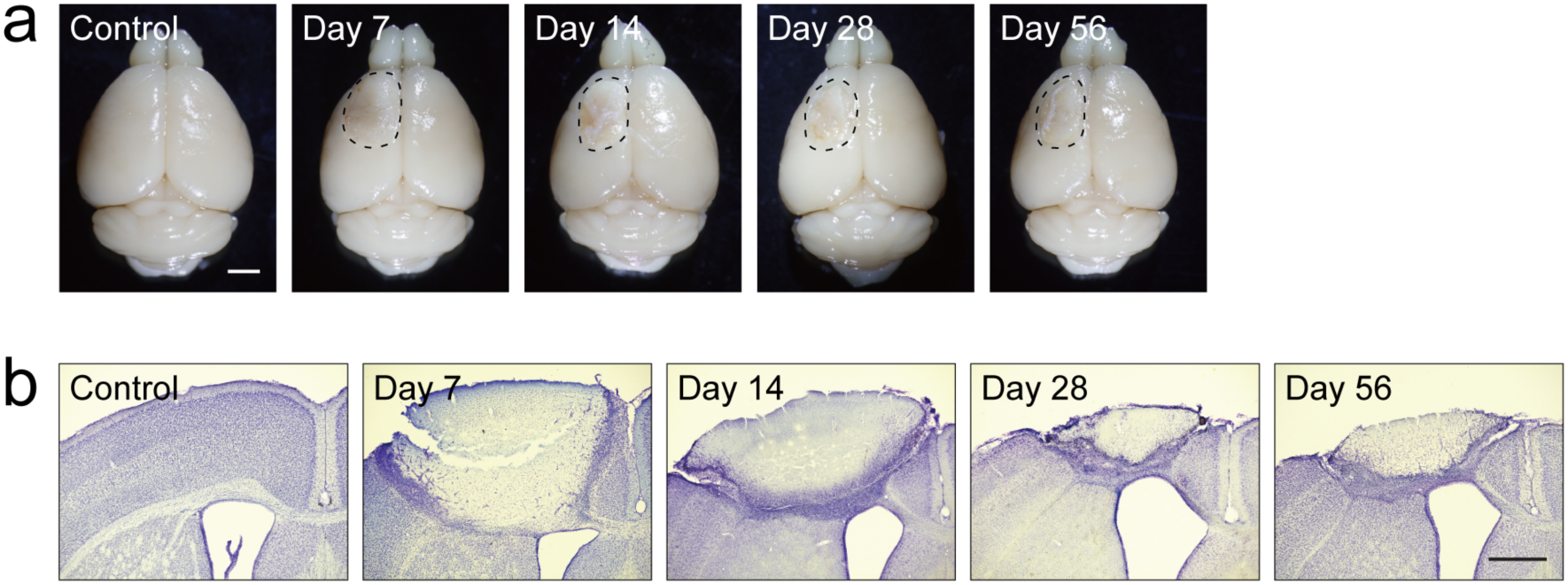
Induction of a large size of stroke in the sensorimotor area. **a**, Representative images of the dorsal view of the brains at days 7–56 after stroke. The lesion areas are indicated by dotted lines. **b**, Representative Nissl staining images of the lesions in the cerebral cortices in coronal sections. Scale bars, 2 mm (**a**); 1 mm (**b**).

**Supplementary Figure 2.**
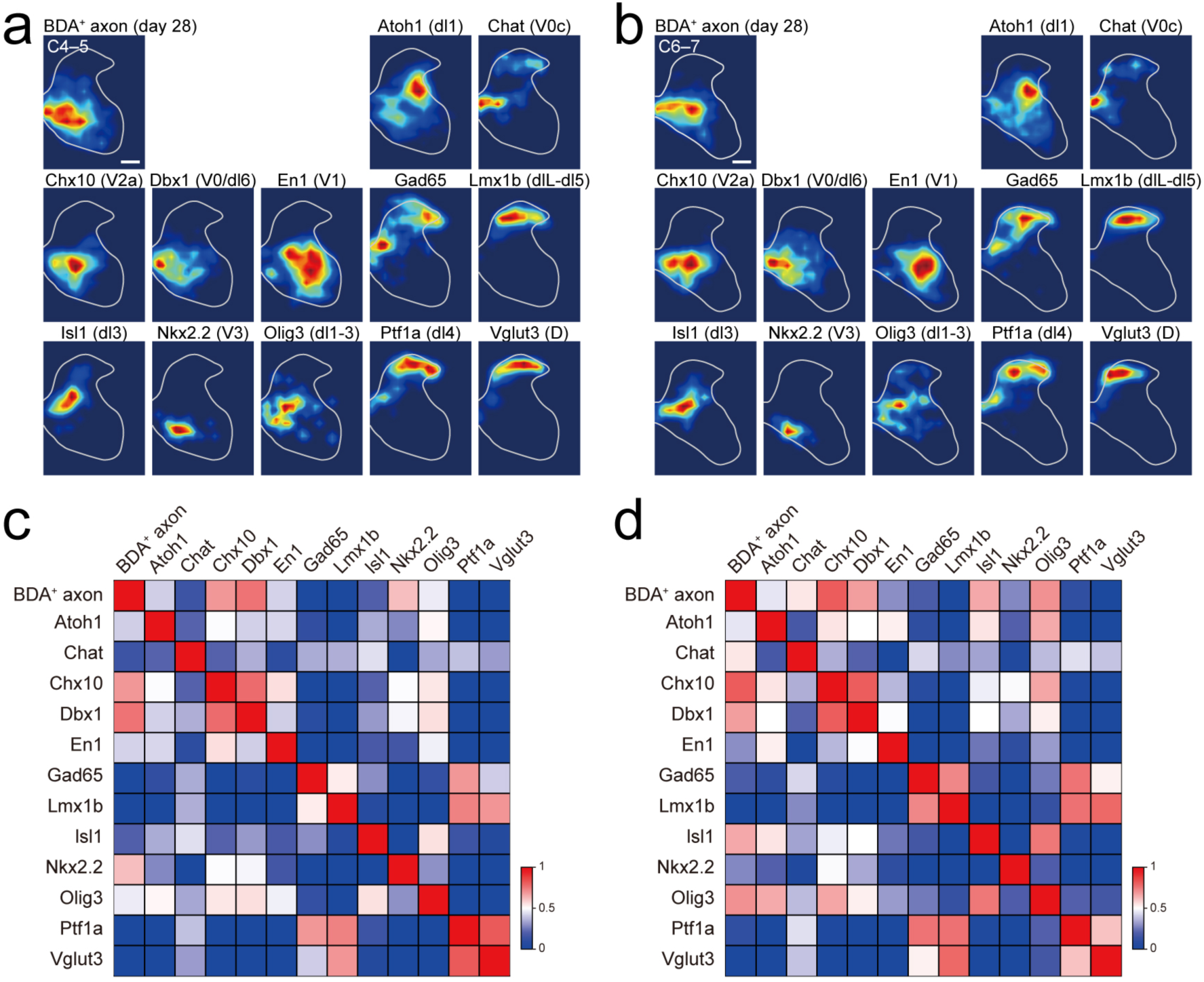
Correlation of spatial distribution of reorganized CST axon projections and spinal interneurons. **a**,**b**, Heatmaps showing the distribution of spared BDA^+^ CST axon projections at day 28 post-stroke and GFP-labeled spinal interneurons at C4–5 (**a**) and C6–7 (**b**). **c**,**d**, Correlation coefficient matrix of the location of CST axons and spinal interneurons at C4–5 (**c**) and C6–7 (**d**). Scale bar, 200 µm.

**Supplementary Figure 3.**
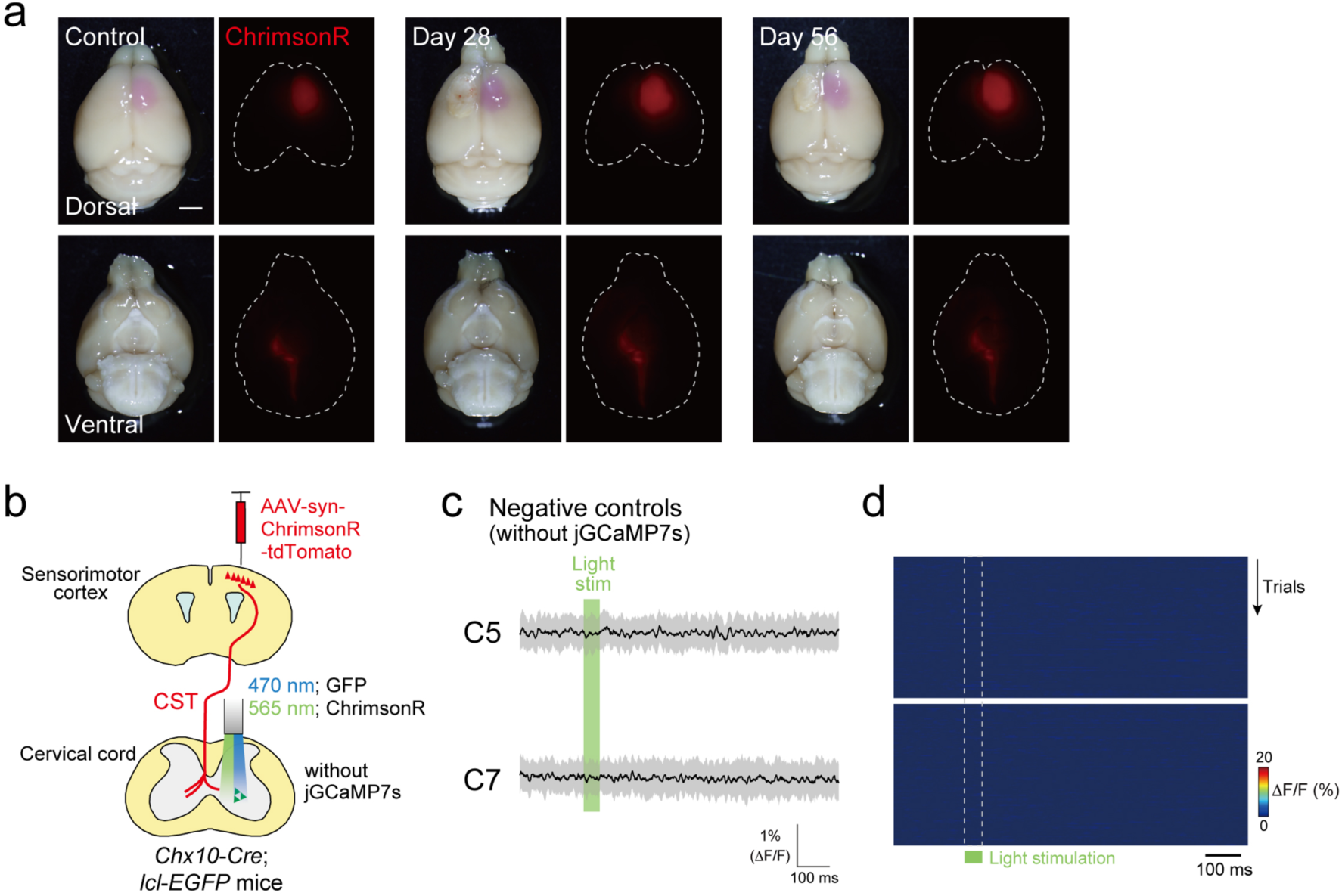
Optical stimulation and fiber photometry in *Chx10-Cre*;*lcl-EGFP* mice. **a**, Representative images of ChrimsonR-tdTomato signals in the brains of control and stroke *Chx10-Cre* mice injected with AAV-Syn-ChrimsonR-tdTomato into the right sensorimotor cortex, which were subjected to the experiments in Fig. 3. Dorsal and ventral views are shown in the top and bottom panels, respectively. Scale bar, 2 mm. **b**, Experimental diagram of fiber photometry with optogenetic stimulation of the CST in negative control *Chx10*-*Cre*;*lcl*-*EGFP* mice without jGCamP7s expression in V2a neurons. **c**, Representative fluorescent responses by optical stimulation at C5 (top) and C7 (bottom) in negative control mice. The solid lines and shaded areas represent the average and standard deviation of fluorescence changes, respectively. Green bar, stimulated period. N = 3 mice (50 trials/one mouse). **d**, Representative kymographs of fluorescence change evoked by optical stimulation at C5 (top) and C7 (bottom) levels (50 trials/one mouse). The green bar and the dotted lines indicate the timing of photostimulation.

**Supplementary Figure 4.**
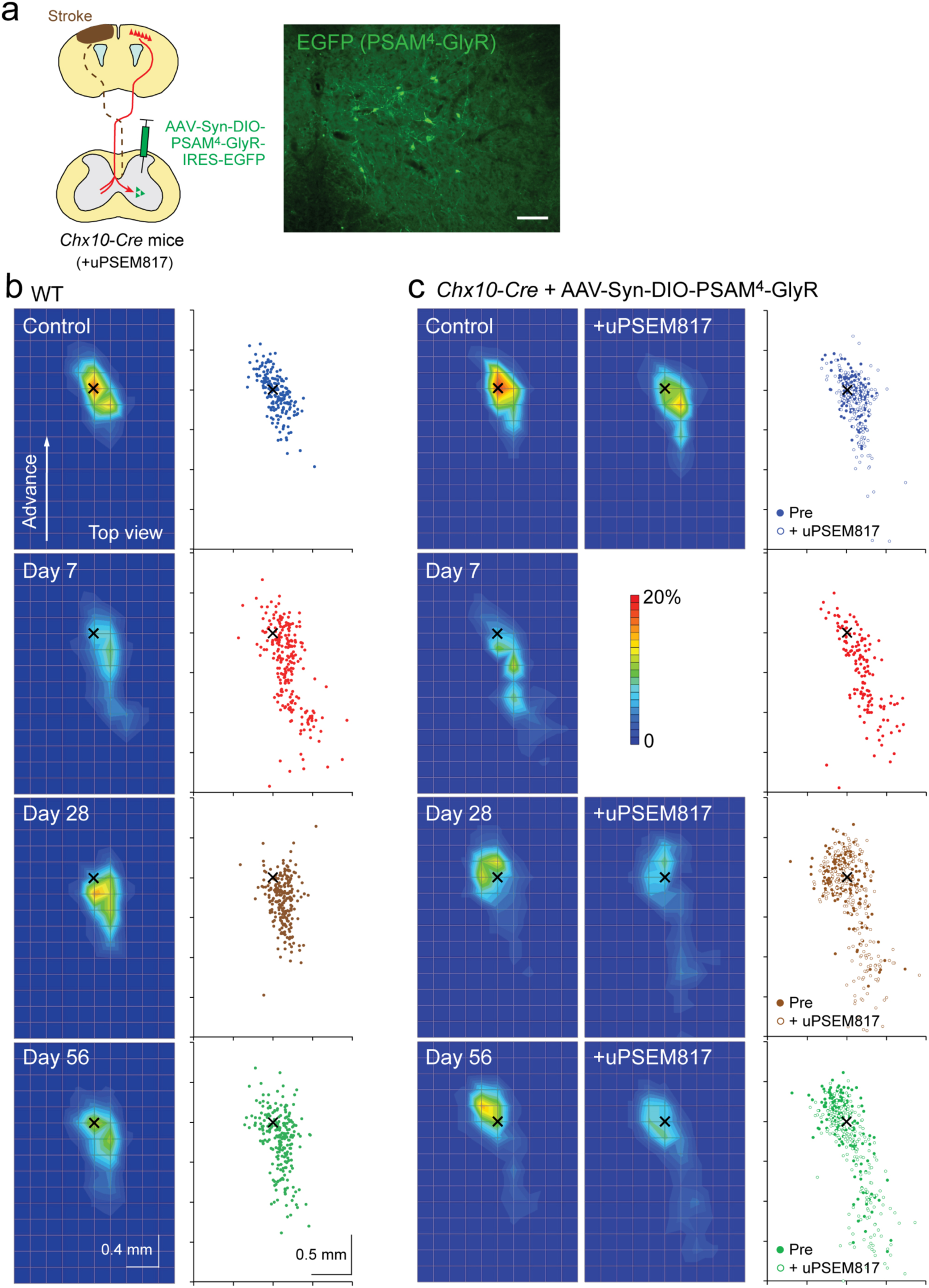
Motor recovery and silencing of V2a interneurons after stroke. **a**, Experimental diagram of AAV-Syn-FLEX-PSAM^4^-GlyR-IRES-EGFP infusion in the cervical cord of *Chx10-Cre* mice and representative image of EGFP (PSAM^4^-GlyR) expression (right, C5 level). Scale bar, 100 µm. **b**, Heatmaps (left, corresponding to Figure 4d) and scatter plots (right) showing the probabilities of the far distal reaching positions of the paw relative to the pellet position (×) in control and post-stroke wild-type (WT) mice. N = 11 mice. **c**, Heatmaps (left, corresponding to Figure 4k) and scatter plots (right) showing the probabilities of the far distal reaching positions of the paw relative to the pellet position (×) in PSAM^4^-GlyR expressing *Chx10- Cre* mice before and after uPSEM817 treatment. N = 6 mice.

**Supplementary Figure 5.**
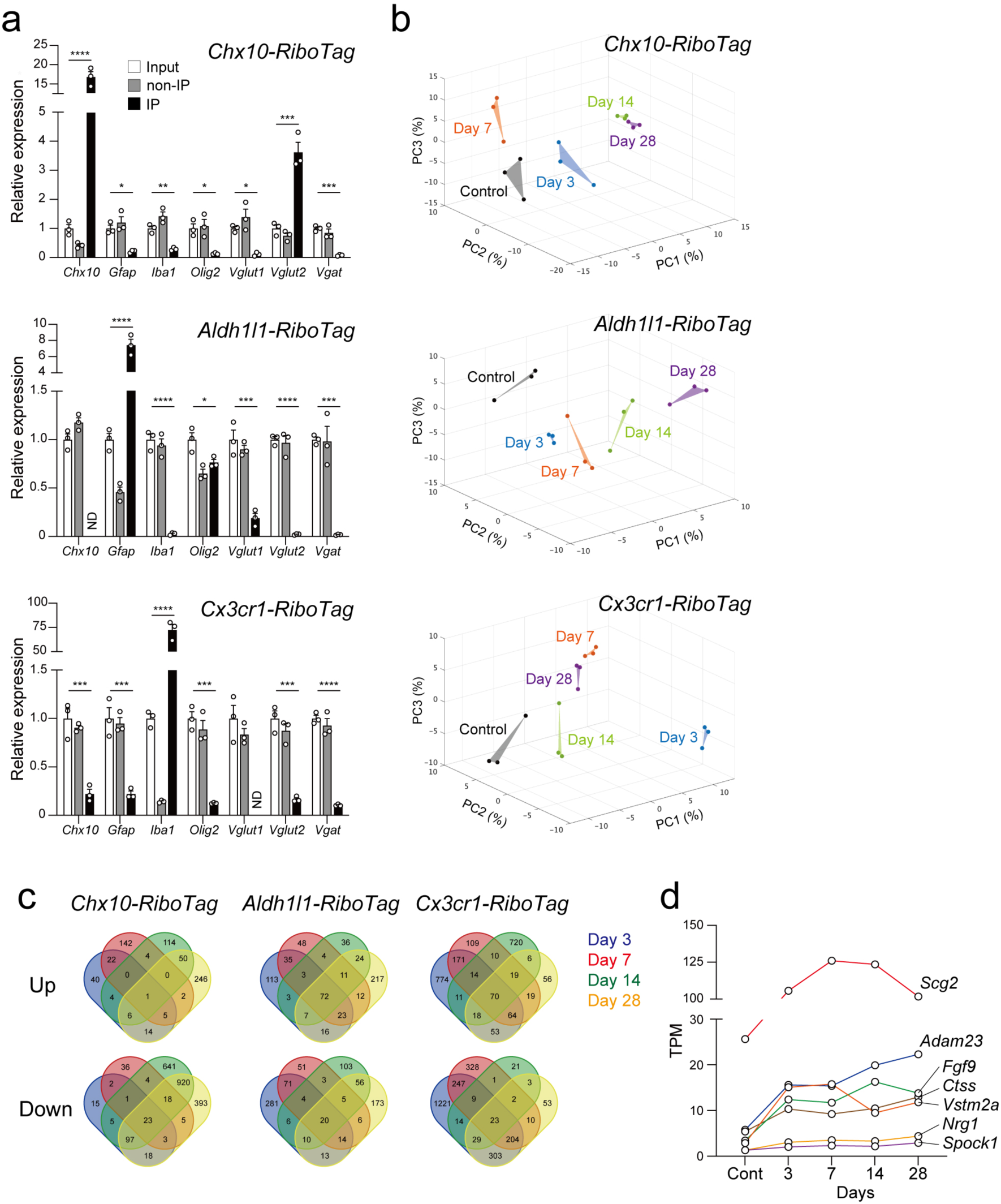
Cell-type specific RiboTag and RNA-seq in the denervated side of the cervical cord after stroke. **a**, Quantitative PCR analysis for evaluating the enrichment of cell-type specific mRNAs. Input, non-immunoprecipitated (non-IP), and IP mRNA were prepared from the right half of the cervical cord (C4–7 level) in intact *Chx10-Cre*;*RiboTag*, *Aldh1l1- CreER*;*RiboTag* and *Cx3cr1-CreER*;*RiboTag* mice. N = 3 mice per group, one-way ANOVA followed by Tukey’s test. \**p* < 0.05, \*\**p* < 0.01, \*\*\**p* < 0.001, \*\*\*\**p* < 0.0001. **b**, PCA analysis of controls and stroke groups of *RiboTag* mice at each time point with the top 1000 most variable genes. **c**, Venn diagrams showing the number of up- and downregulated DEGs in each RiboTag mouse line. **d**, The mean expression level (TPM) of seven constantly upregulated DEGs in astrocytes after stroke (expression data of Fig. 5g).

**Supplementary Figure 6.**
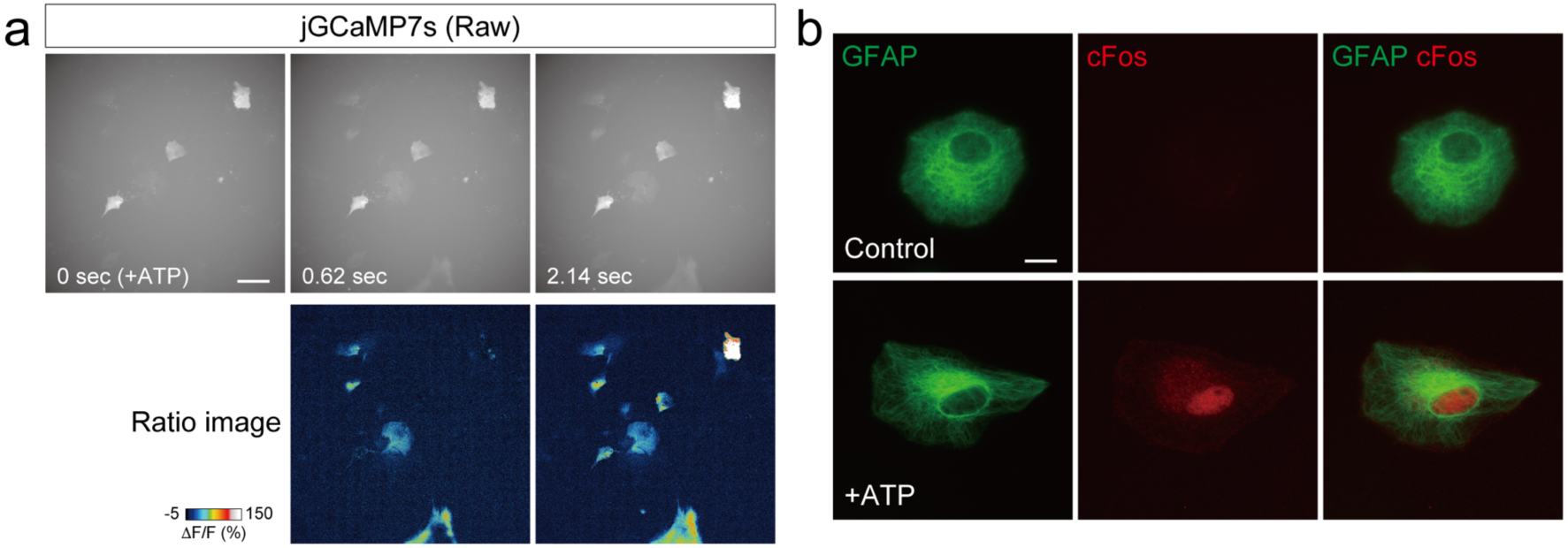
ATP signals induce calcium influx and cFos expression in cultured astrocytes. **a**, Calcium imaging of jGCaMP7s-expressing cultured astrocytes treated with ATP. Raw fluorescence images of jGCaMP7s at pre (0 sec) and post-ATP administration (top) were used for generating the ratio images (bottom). **b**, Representative images of cFos expression (red) induced by ATP stimulation in cultured astrocytes (GFAP, green). Scale bars, 50 µm (**a**) and 10 µm (**b**).

**Supplementary Figure 7.**
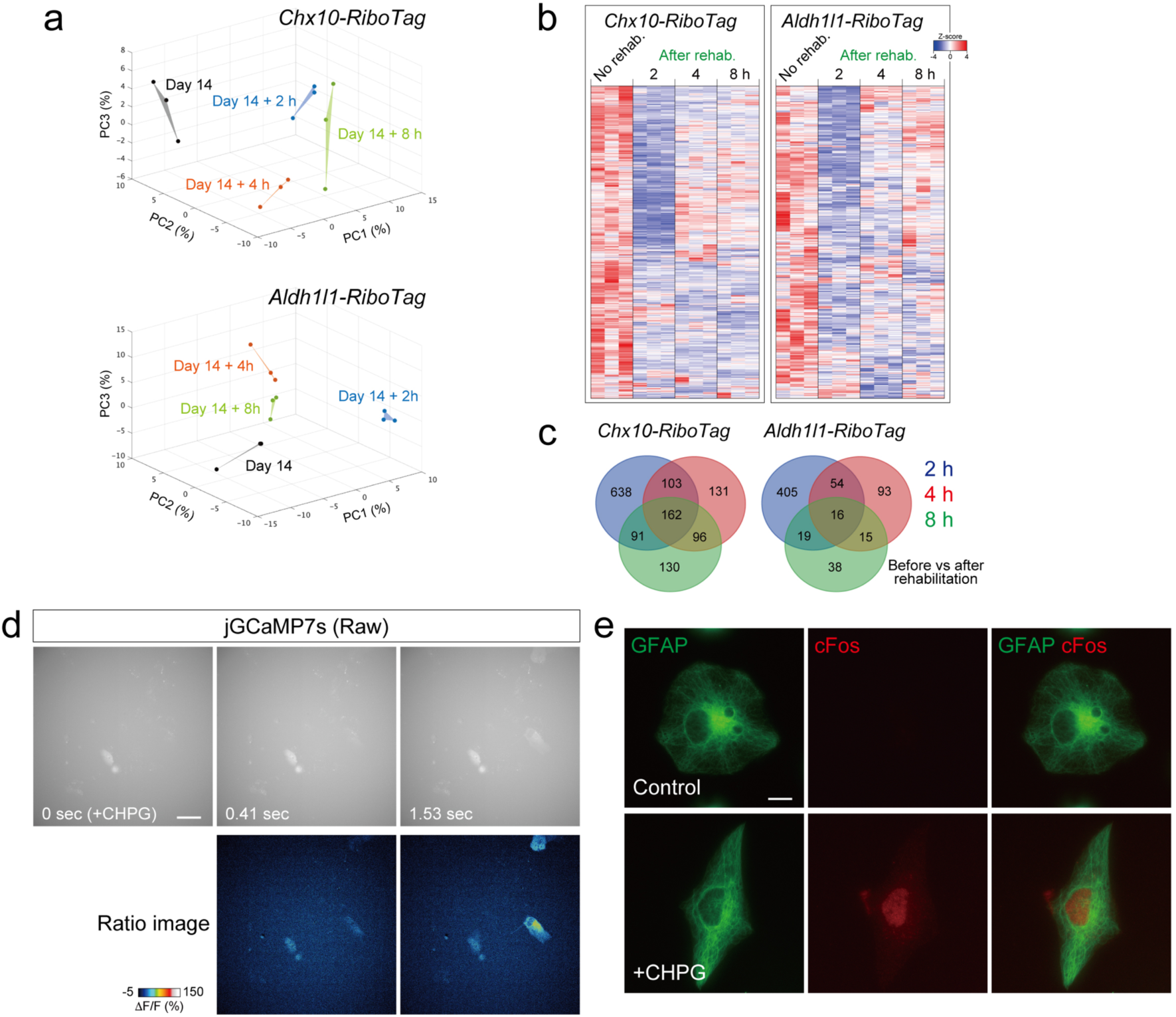
Neuronal activity induced by rotarod rehabilitation leads to gene expression changes in V2a interneurons and astrocytes. **a**, PCA analysis of stroke and rehabilitation groups of *Chx10-Cre*;*RiboTag* and *Aldh1l1-CreER*;*RiboTag* mice at each time point with the top 1000 most variable genes. **b**, Heatmaps showing downregulated DEGs at 2–8 h after rehabilitation in *Chx10-Cre*;*RiboTag* and *Aldh1l1-CreER*;*RiboTag* mice. N = 3 mice for each group. **c**, Venn diagram showing the number of downregulated DEGs in **b**. **d**, Calcium imaging of cultured astrocytes following CHPG treatment. Raw fluorescence images of jGCaMP7s at pre (0 sec) and post-CHPG administration (top) were used for generating the ratio images (bottom). **e**, Representative images of cFos expression (red) by CHPG stimulation in cultured astrocytes (GFAP, green). Scale bars, 50 µm (**d**) and 10 µm (**e**).

**Supplementary Figure 8.**
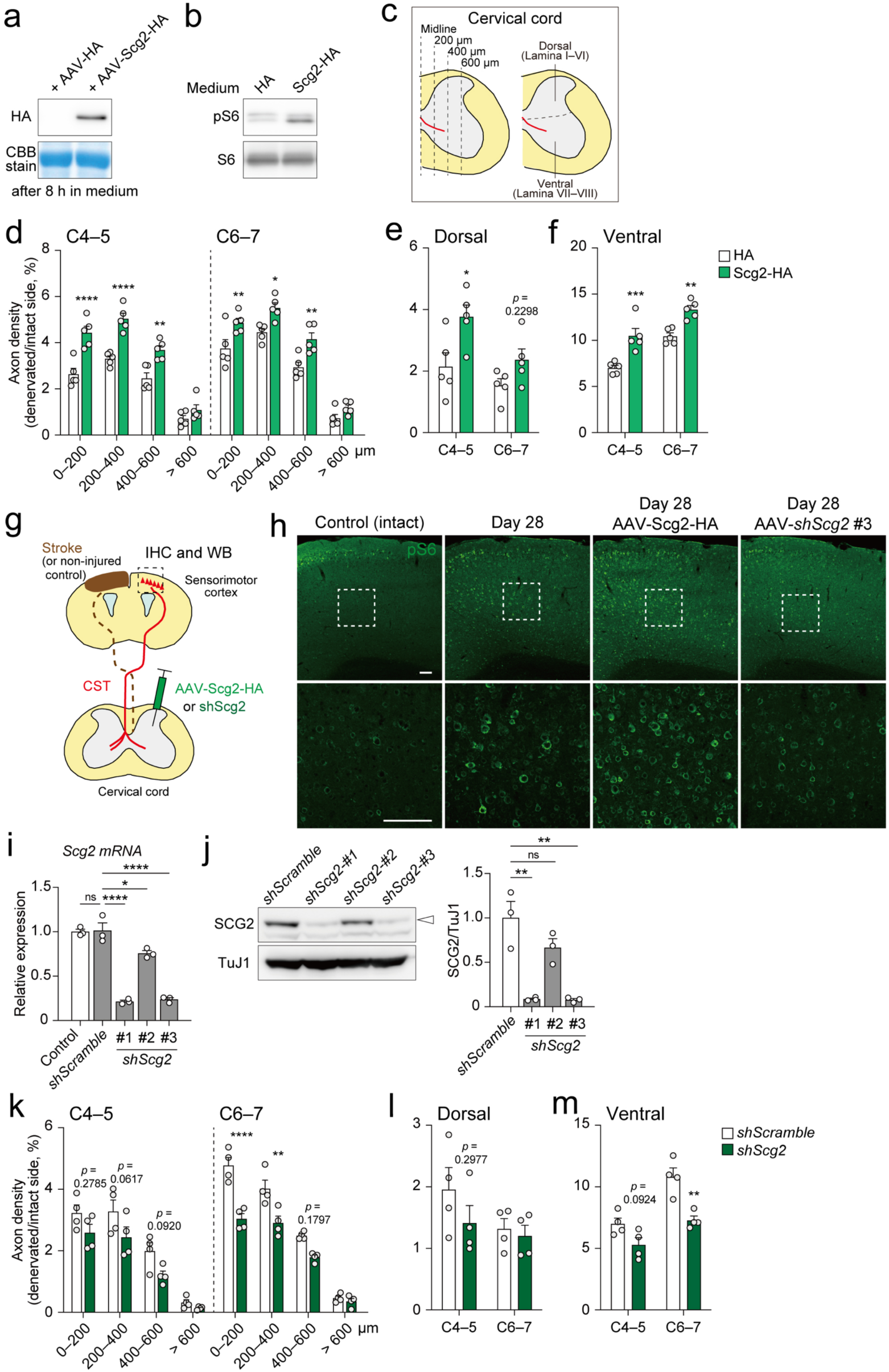
Overexpression and knockdown of *Scg2* change the density of innervating spared CST axons after stroke. **a**, Secretion of SCG2-HA to the conditioned media from Neuro2a cells infected with AAV-CAG-SCG2-HA or control AAV-CAG-HA. **b**, Secreted SCG2-HA in the medium induced phosphorylation of S6 in CGNs. **c**, Schematic representations of mediolateral and dorsoventral areas in the denervated cervical cord for quantification of BDA^+^ axons, segregated with dotted lines. **d**,**e**,**f**, Quantification of CST axon density across the indicated lateral positions (**d**), dorsal (**e**), and ventral laminae (**f**) in the denervated side of C4–5 and C6–7 in control and *Scg2* overexpression group. N = 5, two-way ANOVA followed by Sidak’s post hoc test. **g**, Schematic representations of immunohistochemistry and Western blot analysis for detecting S6 phosphorylation in the contralesional sensorimotor cortex. **h**, Representative confocal images of phospho-S6 (green) in the sensorimotor cortex of control, stroke, and *Scg2* overexpressed/knockdown mice. Lower panels represent high magnification views of the dotted areas in the upper panels. Scale bar, 100 µm. **i**,**j**, Quantification of *Scg2* mRNA in PCR (**i**) and SCG2 proteins in Western blots (**j**) showing the knockdown efficiency of shRNA sequences against *Scg2* in Neuro2a cells transfected with pAAV-CAG-*shScramble* or *shScg2* (#1–#3). N = 3, one-way ANOVA followed by Tukey’s test. **k**,**l**,**m**, Quantification of CST axon density across the indicated lateral positions (**k**), dorsal (**l**), and ventral laminae (**m**) in the denervated side of C4–5 and C6–7 in control *shScramble* and *Scg2* knockdown group. N = 4, two-way ANOVA followed by Sidak’s post hoc test. ns, not significant; \**p* < 0.05, \*\**p* < 0.01, \*\*\**p* < 0.001, \*\*\*\**p* < 0.0001.

**Supplementary Figure 9.**
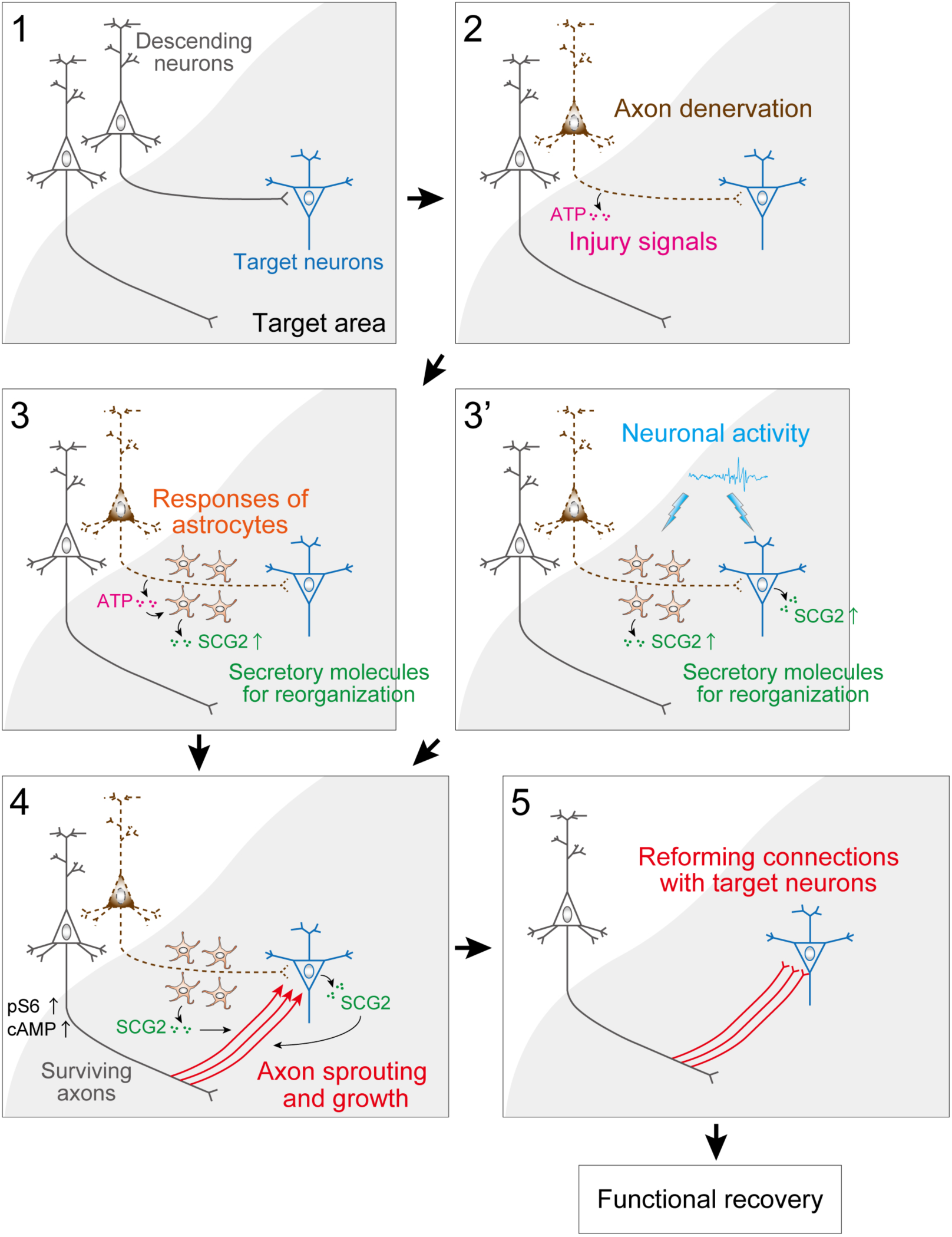
Schematic diagram of the sequential process of neuronal reorganization after CNS injuries. 1) Neural connections of pre- and target post-neurons between distal CNS areas, for example, through the descending pathway such as the corticospinal tract in intact condition. 2) Injuries such as stroke cause loss of neurons and their axons in the distal area, disrupting the connections with the target neurons. Injury signal molecules such as ATP are released from the injured axons. 3) Astrocytes respond to injury signals, which elevate and secrete molecules regulating the reorganization in the denervated area. Our data propose that ATP and Scg2 serve as the injury signal and the master secretory molecule for reorganization, respectively. 3’) In addition to the injury signals, neuronal activity increased such as by rehabilitative training upregulates the expression of Scg2 in astrocytes and target neurons. 4) Surviving axons begin to sprout and elongate in response to the secreted SCG2. pS6 and cAMP levels are elevated in the surviving neurons in response to SCG2. 5) The axons reform connections with the target neurons, and it leads to functional recovery.

Supplementary Table 1. The list of primers and oligonucleotides used in generation of plasmid constructs and quantitative PCR.

Supplementary Video 1. Recovery of single pellet reaching task after stroke in wild-type mice.

Supplementary Video 2. Single pellet reaching task and silencing of V2a neurons after recovery post-stroke in AAV-Syn-FLEX-PSAM^4^-GlyR infused *Chx10*-*Cre* mice.

## Notes

### Competing Interest Statement

The authors have declared no competing interest.

